# Adult organotypic brain slice cultures recapitulate extracellular matrix remodelling in haemorrhagic stroke

**DOI:** 10.1101/2025.07.15.665021

**Authors:** Benjamin J. Hewitt, Lauren Roberts, James A. Roberts, Daniel Fulton, Lisa J. Hill, Phillip Kitchen, Roslyn M. Bill, Hannah F. Botfield

## Abstract

Haemorrhagic stroke is a devastating condition characterised by vessel rupture and free blood within the brain parenchyma or cerebrospinal fluid (CSF) filled spaces. Across the major subtypes of haemorrhagic stroke (subarachnoid, intracerebral, and intraventricular haemorrhages), the presence of blood in the CSF generates significant tissue damage in the first 72 hours after the event, known as early brain injury (EBI). EBI includes neuroinflammation, blood-brain barrier breakdown and dysregulation of extracellular matrix (ECM) dynamics. ECM dysfunction has been shown to trigger fibrosis of the cortical blood vessels, limiting normal CSF circulation and resulting in the buildup of metabolic waste or the development of post-haemorrhagic hydrocephalus. Limiting or preventing this fibrosis may therefore reduce the rate of morbidity experienced by survivors, providing a potential avenue for non-surgical treatment to reduce secondary brain injury post-stroke.

Despite this, current *in vivo* approaches fail to differentiate between the effect of blood products and secondary consequences including intracranial pressure (ICP) elevation and mass effect. Here, we describe an adult rat organotypic brain slice culture (OBSC) model of haemorrhagic stroke which enables the identification of the effect of blood products on ECM dysregulation. We demonstrate the distribution of key cell types across a time course of 0, 3 and 7 days in culture, indicating that such cultures are viable for a minimum of 7 days. Using immunofluorescence staining, Western blotting and RNA sequencing, we show that exposure of OBSCs to lysed blood markedly increases ECM deposition around cortical blood vessels. This is accompanied by dysregulation of ECM regulatory genes and upregulation of inflammation and oxidative stress-related genes, successfully recapitulating the changes seen in human stroke survivors. This versatile *ex vivo* model provides a translational platform to further understanding of haemorrhagic stroke pathophysiology and develop or trial novel therapeutics prior to progression to *in vivo* stroke studies.

## 2. Introduction

Haemorrhagic stroke is a devastating condition characterised by vessel rupture and free blood within the brain parenchyma or cerebrospinal fluid (CSF)-filled spaces, and may be split into intracerebral haemorrhage, subarachnoid haemorrhage (SAH) and intraventricular haemorrhage depending on the location of the bleed. While haemorrhagic stroke only accounts for approximately 20% of stroke cases ^1^, the risk of mortality or profound disability is markedly greater than that posed by ischemic stroke ^2^.

SAH is most commonly triggered by a ruptured aneurysm causing blood to leak into the CSF in the subarachnoid space, with blood also entering the ventricles in higher grade SAH. Early brain injury (EBI) is the injury that occurs in the brain within 72 hours of an SAH and develops due to mechanical factors and the direct effect of free blood. SAH causes an increase in intracranial pressure (ICP), decreased cerebral blood flow (resulting in global ischemia), blood brain barrier (BBB) breakdown, brain oedema, oxidative stress and neuroinflammation, which ultimately leads to neurodegeneration ^3^.

Extracellular matrix (ECM) dynamics in and around the brain are also affected following SAH. There is a fine balance between ECM deposition and degradation which is particularly evident within blood vessel architecture. Under pathological conditions, BBB dysfunction is associated with the degradation of ECM in the basement membrane, mainly through matrix metalloproteinases. This can then release growth factors such as the pleiotropic cytokine transforming growth factor-β1 (TGF-β1) which in turn stimulates vascular smooth muscle cells (SMCs) to promote ECM deposition and repair the BBB ^4^. SAH can result in excessive or dysregulated ECM deposition around major blood vessels on the surface of the brain, contributing to the development of cerebral vasospasm and subsequent delayed cerebral ischemia ^4^. Furthermore, elevated TGF-β1 levels have been implicated in the development of subarachnoid fibrosis and ECM dysfunction in CSF drainage pathways after SAH which can lead to the development of post-haemorrhagic hydrocephalus ^5–8^. Understanding the pathways and mechanisms involved in ECM dysfunction may provide therapeutic targets for reducing further brain injury following SAH.

In both human patients and animal models of SAH, it can be difficult to differentiate between the response of the brain to the mechanical factors that follow a stroke (ICP elevation, mass effect) and the direct reaction of brain tissue to blood products, which is important to understand the pathogenesis of common sequalae to SAH and focus the development of future therapeutics to treat these. In contrast, *ex vivo* models allow us to differentiate between these factors to identity and further understand disease processes. Organotypic brain slices cultures (OBSCs), which were first described in the 1950s for the study of membrane potentials in the cerebral cortex ^9^, have been used in the study of neurodegenerative diseases ^10^, invasive glioma ^11^ and epilepsy ^12^ amongst many other uses. OBSCs provide a three-dimensional tissue architecture and are widely considered to be more representative of the *in vivo* state than cell cultures due to the presence of neurons, glial cells and endothelial cells in a native tissue architecture ^13^. OBSCs are typically prepared from neonatal brain tissues slices as these tend to be more resistant to the mechanical trauma caused by sectioning, making them easier to culture ^14^. However, these neonatal models are unlikely to recapitulate the complexity or maturity of adult tissues, as is shown by different responses to sectioning. For this reason, adult OBSC models are essential in modelling a disease which primarily affects middle to older-aged adults ^15^. Whilst adult rodent OBSCs are now well established, there has been very limited prior exploration of their application to haemorrhagic stroke research.

Attempts at short-term (minutes to hours) haemorrhagic stroke brain slice models have been reported with a focus on neuronal survival, though it is not clear whether they were from adult or neonatal animals ^16^. Therefore, the development and validation of an adult OBSC system capable of modelling haemorrhagic stroke remains an important challenge.

Here, we describe the generation and validation of OBSCs derived from healthy adult rats and their application to the *ex vivo* modelling of post-haemorrhagic stroke ECM dysfunction and inflammation. *Ex vivo* recapitulation of ECM dynamics may provide an opportunity for furthering understanding of the molecular mechanisms underlying this key factor in post- stroke brain injury and offer a platform in which to trial novel therapeutics prior to use *in vivo*.

## 3. Methods

### 3.1. Tissue culture

All tissue culture was performed under aseptic conditions using a class II microbiological safety cabinet (Monmouth Guardian T1200). Tissues were cultured at 37°C, 5% CO2 and 100% relative humidity.

Dissection media for OBSC generation comprised Earle’s balanced salt solution supplemented to a final concentration of 25 mM HEPES. OBSC culture media comprised 1:1 Dulbecco’s Modified Eagle Media (DMEM):Neurobasal A, supplemented with final concentrations of 2% (v/v) B-27, 2 mM L-glutamine and 1% penicillin/streptomycin solution.

### 3.2. Experimental animals

Male Sprague-Dawley rats were group housed with access to food and water *ad libitum*, on a 12-hour light/dark cycle. All studies were conducted at the University of Birmingham, and both the establishment and investigators held relevant UK Home Office licenses. Animals were monitored daily and group housed, with no regulated procedures performed. Animals were euthanized according to Schedule 1 of the Animals (Scientific Procedures) Act 1986 at a weight of 130-150 g by intraperitoneal injection of 200 mg/ml pentobarbitol, 4 ml/kg and decapitated following cessation of circulation. Blood was captured in a sterile, un-heparinised 50 ml centrifuge tube and stored at -20°C until required.

### 3.3. Organotypic brain slice culture (OBSC) generation

The brain was rapidly removed from the skull and the cerebellum and olfactory bulbs excised with a sterile scalpel. A flat face was produced on the rostral and caudal edges of the brain by cutting (Figure 1), and the brain adhered to the chuck of a vibratome (7000smz, Campden Instruments) (caudal side down) with cyanoacrylate glue. The brain was then mounted into the vibratome bath, containing 250 ml dissection medium on ice, bubbled with medical oxygen at 2 psi. 250 µM sections were produced with the vibratome set to a 1.25 mm amplitude at 80 Hz and sections stored in the dissection media bath until completion.

**Figure 1.**
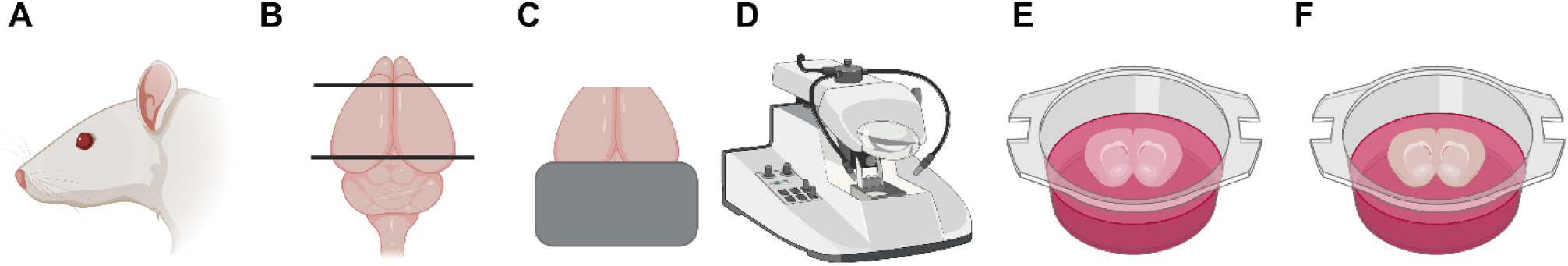
Steps in the generation of adult rat brain OBSCs. A) Sacrifice and brain removal, B) Removal of olfactory bulbs and cerebellum, including production of a flat edge on the caudal aspect of the cerebrum, C) Mounting of brain (caudal face down) onto vibratome chuck with cyanoacrylate glue, D) Vibratome slicing of brain (250 µM sections, 80 Hz, 1.25 mm amplitude), E) Brain slices added to 6-well cell culture inserts, F) Media removed to bring brain slice to air-liquid interface.

Slices were then transferred to 6-well polyethylene culture inserts (657641, Greiner) using a fine paintbrush and spatula. To facilitate this, 1 ml of culture medium was initially added to each insert to enable floating of the slice, which was removed once the slice was correctly positioned. Culture inserts were added to a standard 6-well plate (3516, Corning), and 1.5 ml of OBSC culture medium added to the bottom well with the top insert containing the brain slice left dry. Slices were then either fixed immediately or on day in-vitro (DIV) 3 or 7 using 4% paraformaldehyde (PFA) in PBS for one hour at room temperature, followed by two, one-hour phosphate buffered saline (PBS) washes at room temperature prior to storage in PBS at 4°C.

### 3.4. Blood-treated OBSCs

OBSCs destined for blood treatment as a model of haemorrhagic stroke were generated as described above and exposed to blood immediately or after 7 days of pre-culture.

Blood from all animals was pooled, vortexed and shaken vigorously to break up clots, and diluted 1:1 with PBS. 6, 24 or 48 hours prior to the experiment endpoint, 175 µl of diluted blood (or PBS control) was added to the wells assigned for blood treatment after removal of an equal volume of medium. Brain slices for immunohistochemical analysis were then fixed as previously described, prior to washing through two changes of PBS and storage at 4°C.

Slices destined for protein or RNA extraction were dissected under a surgical microscope, with the cortex removed (taking care to avoid capturing any tissue from the corpus callosum), snap frozen in liquid nitrogen and stored at -80°C.

### 3.5. Immunohistochemical staining and imaging

OBSCs were divided into two hemispheres prior to staining. A full protocol for staining and clearing of OSBCs can be found in the supplemental materials (‘Staining and clearing of OBSCs’). Antibodies were diluted in PBS with 0.2% (v/v) triton X-100, 6% (v/v) goat serum and 20% (v/v) DMSO, with 0.2 ml of diluted antibody added to each slice in a 24-well plate. Imaging was performed using an AxioScan Z.1 (Zeiss) for blood exposed sections and an AxioScan 7 (Zeiss) for timepoint studies maintaining the same imaging settings across all slides within an analysis group. 4,6-diamidino-2-phenylindole (DAPI) was included at a 1:5000 dilution of a 5 mg/ml stock in PBS.

**Table 1.** Antibody dilutions used in immunofluorescent staining of OBSCs.

| <u>Antibody</u> | <u>Dilution</u> |
| --- | --- |
| Anti-rat endothelial cell antigen 1 (RECA-1), mouse | 1:100 |
| Anti-fibronectin, rabbit | 1:400 |
| Anti-alpha smooth muscle actin (αSMA), mouse | 1:500 |
| Anti-cleaved caspase 3 (CCP3), rabbit | 1:500 |
| Anti-NeuN, mouse | 1:500 |
| Anti-GFAP, mouse | 1:400 |
| Anti-collagen IV, rabbit | 1:500 |
| Anti-IBA1, rabbit | 1:500 |
| AF488 conjugated anti-rabbit IgG, highly cross-absorbed | 1:1000 |
| AF594 conjugated anti-mouse IgG, highly cross-absorbed | 1:1000 |

### 3.6. Image processing and analysis

Image processing was performed using FIJI ^17^, with the operator blinded to the treatment or culture conditions of each image by displaying only numerical filenames. Representative images shown have been enhanced for display by modification of window and level, performed uniformly across all images within a comparison group or stain. No image enhancement was performed on images prior to image analysis.

Images were processed using a FIJI macro ^17^, made available both on GitHub (https://github.com/Dr-ben-hewitt/OBSC-quantification-macros) and in the supplementary materials. In brief, this performed a maximum intensity projection and presented the user with an image in the red channel only (corresponding to blood vessel staining with RECA-1). Five regions of interest (ROIs) were then manually selected in the cortex of each slice and processed for background subtraction (100 pixel radius for RECA-1 channel, 1000 pixel radius for fibronectin and collagen IV), followed by automatic thresholding of blood vessels (RenyiEntropy method) and ECM proteins (Triangle method), with image calculation then performed to find the average of these two sets of features.

Mean pixel intensity per unit area was calculated by user selection of 3 ROIs per image (with the user shown only the DAPI channel to limit bias), followed by background subtraction (50 pixel radius) in FIJI, automated measurement of the integrated ROI pixel intensity and division of this value by the area of the ROI.

### 3.7. Protein extraction and Western blotting

Protein extraction was performed by mechanical homogenisation in 250 µl radioimmunoprecipitation assay (RIPA) buffer, supplemented with protease and phosphatase inhibitors per manufacturers guidance. Samples were then incubated on a roller at 4°C for 2 hours and centrifuged at 10,000×*g* for 20 minutes at 4°C. Supernatant was then harvested, aliquoted and stored at -80°C until required. Protein concentration was assessed by BCA assay, following the manufacturer’s instructions. SDS-PAGE was performed using 2.5 µg of protein in a 20 µl volume, containing 5 µl LDS sample buffer and 2 µl reducing agent, on 4- 12% bis-tris polyacrylamide gels using MOPS-SDS running buffer. Electrophoresis was performed at 120 V for 90 minutes. Proteins were then transferred to a nitrocellulose membrane using NuPAGE transfer buffer at 25 V for 1 hour. Blocking was performed by immersion in 5% (w/v) skimmed milk in tris-buffered saline with 0.1% (v/v) Tween-20 (TBST) for one hour at room temperature. Blots were then incubated in primary antibodies for fibronectin or collagen IV and alpha-actin diluted in blocking buffer for one hour at room temperature, washed three times in TBST (15 minutes each, room temperature), and incubated in secondary antibody (1:5000 anti-IgG HRP conjugated) for one hour at room temperature. Blots were then washed 3x in TBST (10 minutes each, room temperature) prior to immersion in chemiluminescent substrate for 5 minutes (room temperature) and imaging (G:Box chemi XRQ, Syngene).

### 3.8. RNA extraction, sequencing and RT-qPCR

Samples were mechanically disrupted and homogenised in 1 ml TRIzol and 200 µl chloroform was then added. Samples were then mixed by shaking and centrifuged at 12,000\**g* for 15 minutes at 4°C. The upper aqueous phase was removed and transferred to a new tube, and an equal volume of molecular grade 70% ethanol added. RNA was then extracted from the sample with an RNEasy mini extraction kit (Qiagen), following the manufacturer’s instructions for a 40 µl elution volume in RNase free water. RNA was then quantified by spectrophotometery (N50 NanoPhotometer, Implen). Sample quality control (integrity, purity and concentration via Agilent 5400) and paired-end mRNA sequencing was performed by a contracted 3^rd^ party (Novogene, UK) using a NovaSeq X Plus (Illumina) at a read depth of 20 million reads per sample.

Alignment against a Sprague Dawley rat genome (rn_celera) was performed by Novogene, followed by mapping to the same reference genome using the featureCounts tool ^18^ on the Galaxy platform ^19^, followed by quality control using MultiQC ^20^. Analysis of differentially expressed genes was performed with DESeq2 ^21^, with comparisons made of expression in blood-treated samples against an untreated control. Differentially expressed genes were defined as those with an adjusted p-value (*pAdj)* < 0.05, and a fold change cutoff of 2 was used for display of DEGs.

Gene ontology (GO) analysis was performed on all differentially expressed genes (with no fold-change cutoff) using goseq ^22^, with gene categories retrieved from the rn6 assembly due to unavailability of gene categories for the rn_celera assembly. All visualisation of RNA-sequencing data was performed using GraphPad Prism 10.3.

Validation of the top 6 most upregulated and top 6 most downregulated genes was performed using RT-qPCR. In brief, cDNA conversion was performed with a Tetro cDNA kit using Oligo(dT)18 primers and 100 ng of RNA. cDNA was then further diluted by addition of 20 μl molecular biology grade water. RT-qPCR mastermix comprised 100 μl SYBR green, 78 μl water and 1 μl of each forward and reverse primer (reconstituted to 100 μM). 9 μl mastermix was added to each well of a 384-well PCR plate, followed by 1 μl cDNA. RT-qPCR was performed using a QuantStudio 5, with reaction conditions shown in supplemental materials **Error! Reference source not found.**1. Oligonucleotide primers used are shown in supplemental materials Table S2

### 3.9. Statistical analysis

All statistical analysis was performed in GraphPad Prism 10.3, with a significance level of 0.05. Statistical tests are described in the relevant figure legends.

RT-qPCR data were analysed with QBase+ (Biogazelle, Belgium) using the calibrated normalised relative quantity (CNRQ) method. Four housekeeping genes were assessed using the geNorm algorithm within QBase+ and the two housekeeping genes with lowest mean variability were used as reference genes for sample normalisation.

### 3.10. Key resources table

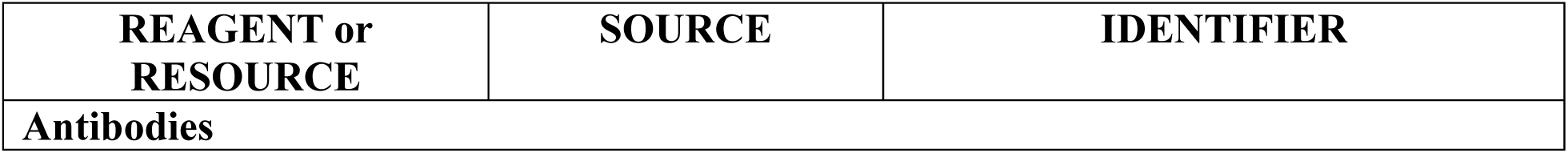
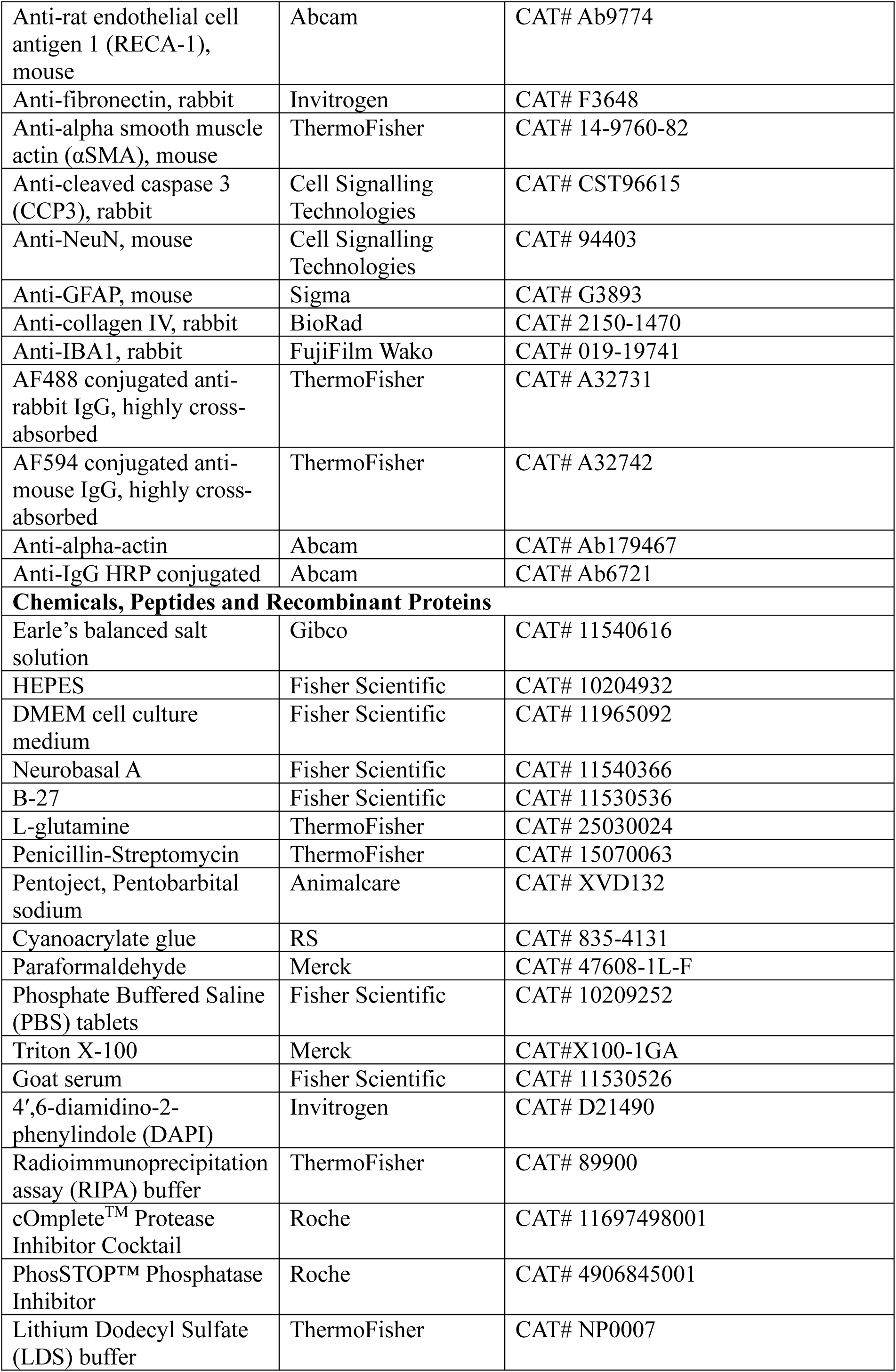
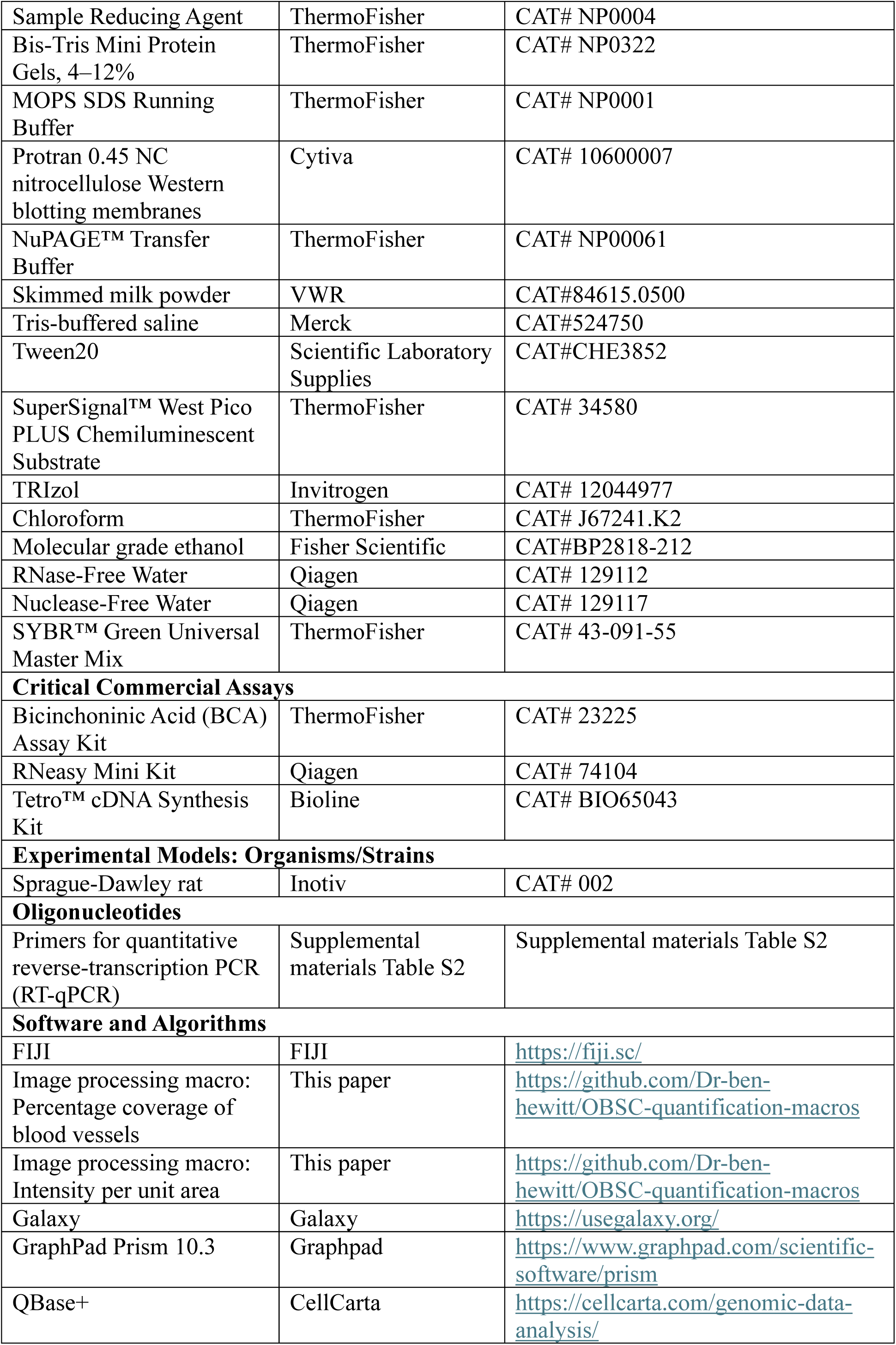
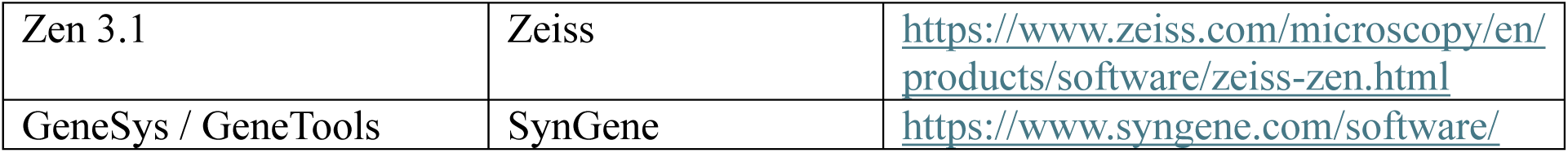

| REAGENT or RESOURCE | SOURCE | IDENTIFIER |
| --- | --- | --- |
| Antibodies |  |  |
| **Running title:** Modelling ECM dysfunction in haemorrhagic stroke |  |  |
| Anti-rat endothelial cell antigen 1 (RECA-1), mouse | Abcam | CAT# Ab9774 |
| Anti-fibronectin, rabbit | Invitrogen | CAT# F3648 |
| Anti-alpha smooth muscle actin (αSMA), mouse | ThermoFisher | CAT# 14-9760-82 |
| Anti-cleaved caspase 3 (CCP3), rabbit | Cell Signalling Technologies | CAT# CST96615 |
| Anti-NeuN, mouse | Cell Signalling Technologies | CAT# 94403 |
| Anti-GFAP, mouse | Sigma | CAT# G3893 |
| Anti-collagen IV, rabbit | BioRad | CAT# 2150-1470 |
| Anti-IBA1, rabbit | FujiFilm Wako | CAT# 019-19741 |
| AF488 conjugated anti-rabbit IgG, highly cross-absorbed | ThermoFisher | CAT# A32731 |
| AF594 conjugated anti-mouse IgG, highly cross-absorbed | ThermoFisher | CAT# A32742 |
| Anti-alpha-actin | Abcam | CAT# Ab179467 |
| Anti-IgG HRP conjugated | Abcam | CAT# Ab6721 |
| **Chemicals, Peptides and Recombinant Proteins** |  |  |
| Earle's balanced salt solution | Gibco | CAT# 11540616 |
| HEPES | Fisher Scientific | CAT# 10204932 |
| DMEM cell culture medium | Fisher Scientific | CAT# 11965092 |
| Neurobasal A | Fisher Scientific | CAT# 11540366 |
| B-27 | Fisher Scientific | CAT# 11530536 |
| L-glutamine | ThermoFisher | CAT# 25030024 |
| Penicillin-Streptomycin | ThermoFisher | CAT# 15070063 |
| Pentoject, Pentobarbital sodium | Animalcare | CAT# XVD132 |
| Cyanoacrylate glue | RS | CAT# 835-4131 |
| Paraformaldehyde | Merck | CAT# 47608-1L-F |
| Phosphate Buffered Saline (PBS) tablets | Fisher Scientific | CAT# 10209252 |
| Triton X-100 | Merck | CAT#X100-1GA |
| Goat serum | Fisher Scientific | CAT# 11530526 |
| 4',6-diamidino-2-phenylindole (DAPI) | Invitrogen | CAT# D21490 |
| Radioimmunoprecipitation assay (RIPA) buffer | ThermoFisher | CAT# 89900 |
| cOmplete<sup>TM</sup> Protease Inhibitor Cocktail | Roche | CAT# 11697498001 |
| PhosSTOP<sup>TM</sup> Phosphatase Inhibitor | Roche | CAT# 4906845001 |
| Lithium Dodecyl Sulfate (LDS) buffer | ThermoFisher | CAT# NP0007 |
| **Running title:** Modelling ECM dysfunction in haemorrhagic stroke |  |  |
| Sample Reducing Agent | ThermoFisher | CAT# NP0004 |
| Bis-Tris Mini Protein Gels, 4–12% | ThermoFisher | CAT# NP0322 |
| MOPS SDS Running Buffer | ThermoFisher | CAT# NP0001 |
| Protran 0.45 NC nitrocellulose Western blotting membranes | Cytiva | CAT# 10600007 |
| NuPAGE™ Transfer Buffer | ThermoFisher | CAT# NP00061 |
| Skimmed milk powder | VWR | CAT#84615.0500 |
| Tris-buffered saline | Merck | CAT#524750 |
| Tween20 | Scientific Laboratory Supplies | CAT#CHE3852 |
| SuperSignal™ West Pico PLUS Chemiluminescent Substrate | ThermoFisher | CAT# 34580 |
| TRIzol | Invitrogen | CAT# 12044977 |
| Chloroform | ThermoFisher | CAT# J67241.K2 |
| Molecular grade ethanol | Fisher Scientific | CAT#BP2818-212 |
| RNase-Free Water | Qiagen | CAT# 129112 |
| Nuclease-Free Water | Qiagen | CAT# 129117 |
| SYBR™ Green Universal Master Mix | ThermoFisher | CAT# 43-091-55 |
| **Critical Commercial Assays** |  |  |
| Bicinchoninic Acid (BCA) Assay Kit | ThermoFisher | CAT# 23225 |
| RNeasy Mini Kit | Qiagen | CAT# 74104 |
| Tetro™ cDNA Synthesis Kit | Bioline | CAT# BIO65043 |
| **Experimental Models: Organisms/Strains** |  |  |
| Sprague-Dawley rat | Inotiv | CAT# 002 |
| **Oligonucleotides** |  |  |
| Primers for quantitative reverse-transcription PCR (RT-qPCR) | Supplemental materials Table S2 | Supplemental materials Table S2 |
| **Software and Algorithms** |  |  |
| FIJI | FIJI | https://fiji.sc/ |
| Image processing macro: Percentage coverage of blood vessels | This paper | https://github.com/Dr-ben-hewitt/OBSC-quantification-macros |
| Image processing macro: Intensity per unit area | This paper | https://github.com/Dr-ben-hewitt/OBSC-quantification-macros |
| Galaxy | Galaxy | https://usegalaxy.org/ |
| GraphPad Prism 10.3 | Graphpad | https://www.graphpad.com/scientific-software/prism |
| QBase+ | CellCarta | https://cellcarta.com/genomic-data-analysis/ |
| Zen 3.1 | Zeiss | https://www.zeiss.com/microscopy/en/products/software/zeiss-zen.html |
| GeneSys / GeneTools | SynGene | https://www.syngene.com/software/ |

## 4. Results

### 4.1. Health of the OBSCs after 7 days in culture

Cleaved caspase-3 (CPP3) staining was used as a marker of apoptosis. Overall, there was an increase in CCP3-positive staining from DIV0 to DIV7 in the cortex (Figure 2A) and other areas of the OBSC (Figure S3). Quantification of the mean CCP3-positive pixel intensity per unit area revealed a significant increase in CCP3 signal in the cortex (Figure 2B), increasing from 1.21×10^-6^ at 0 days to 2.04×10^-6^ at 3 days (*P* = 0.0081) and 2.72×10^-6^ at 7 days (*P* = 0.0001). A significant increase in CCP3 signal between DIV0 and DIV7 was also noted in the basal ganglia (Figure S3).

**Figure 2.**
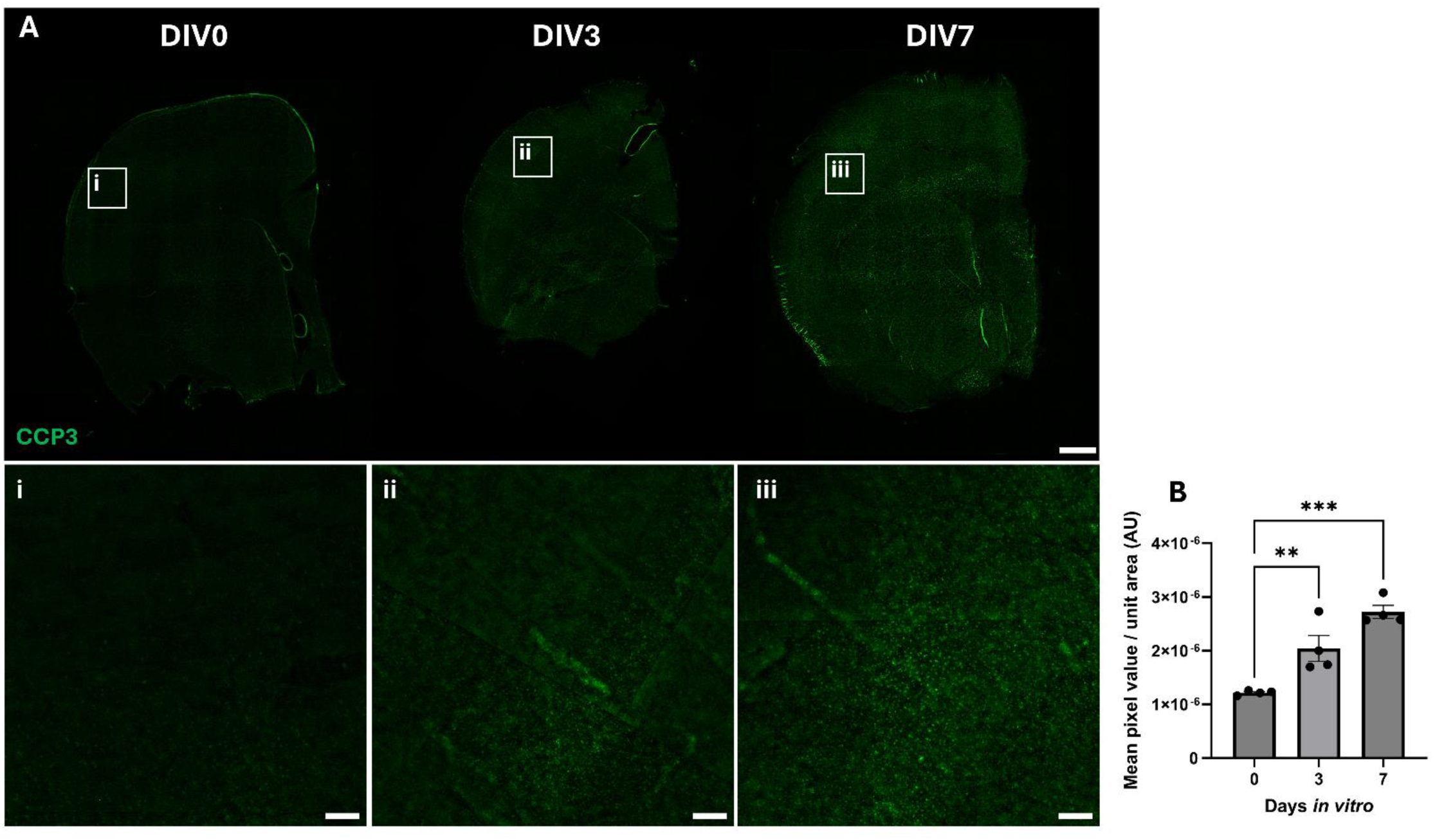
Distribution and intensity of CCP3 throughout control and cultured OBSCs. A) Representative immunofluorescent staining of OBSCs (CCP3, green and DAPI, blue) at 0 3 and 7 days *in vitro* (scale bar = 1,000 µm). Magnified regions of the cortex of section are shown in insets ‘i’, ‘ii’ and ‘iii’ for DIV0, 3 and 7 respectively (scale bar = 100 µm for all). Quantification of mean CCP3-stained pixel intensity per unit area in the cortex was also performed (B). Each image quantification datapoint is mean of 6 regions of interest from 2 brain slices per animal (3 regions per slice), ± SEM. One-way ANOVA with Dunnett’s multiple comparisons test (all to all). ns = not significant, ** *P* < 0.01, *** *P* < 0.001. *N* = 4 animals, *α* = 0.05.

### 4.2. Alterations in the organotypic cytoarchitecture of the cortex after 7 days in culture

Previous studies suggested OBSCs need to be cultured for 7-14 days prior to use in order to allow the slices to recover from the trauma of vibratome sectioning. Therefore, the cellular composition of OBSCs was assessed at DIV0, DIV3 and DIV7.

NeuN, a neuronal cell marker, demonstrated stable levels in the cortex of OBSCs at DIV0, DIV3 and DIV7 (Figure 3A), with no significant differences observed in NeuN staining intensity between DIV0 (mean pixel intensity per unit area of 1.21x10^-5^) and DIV3 or 7 (8.49x10^-6^ and 6.24x10^-6^ respectively, Figure 3B). In the cortex, glial fibrillary acidic protein (GFAP)-positive astrocytes (Figure 3C) appeared to show a marked reduction in staining from DIV0 to DIV7, particularly with a loss in the glia limitans. This was further supported by a significant reduction in quantified GFAP mean pixel value at both DIV3 and DIV7 compared to DIV0. Mean pixel intensity per area of GFAP was reduced by more than 50% from 5.3×10^-6^ at DIV0 to 2.0×10^-6^ and 2.5×10^-6^ at DIV3 and DIV7 (*P* < 0.0001 and *P* = 0.0001, respectively) (Figure 3D). In contrast, GFAP-positive astrocytes were maintained within the deeper subcortical white matter (supplemental materials, Figure S4).

**Figure 3.**
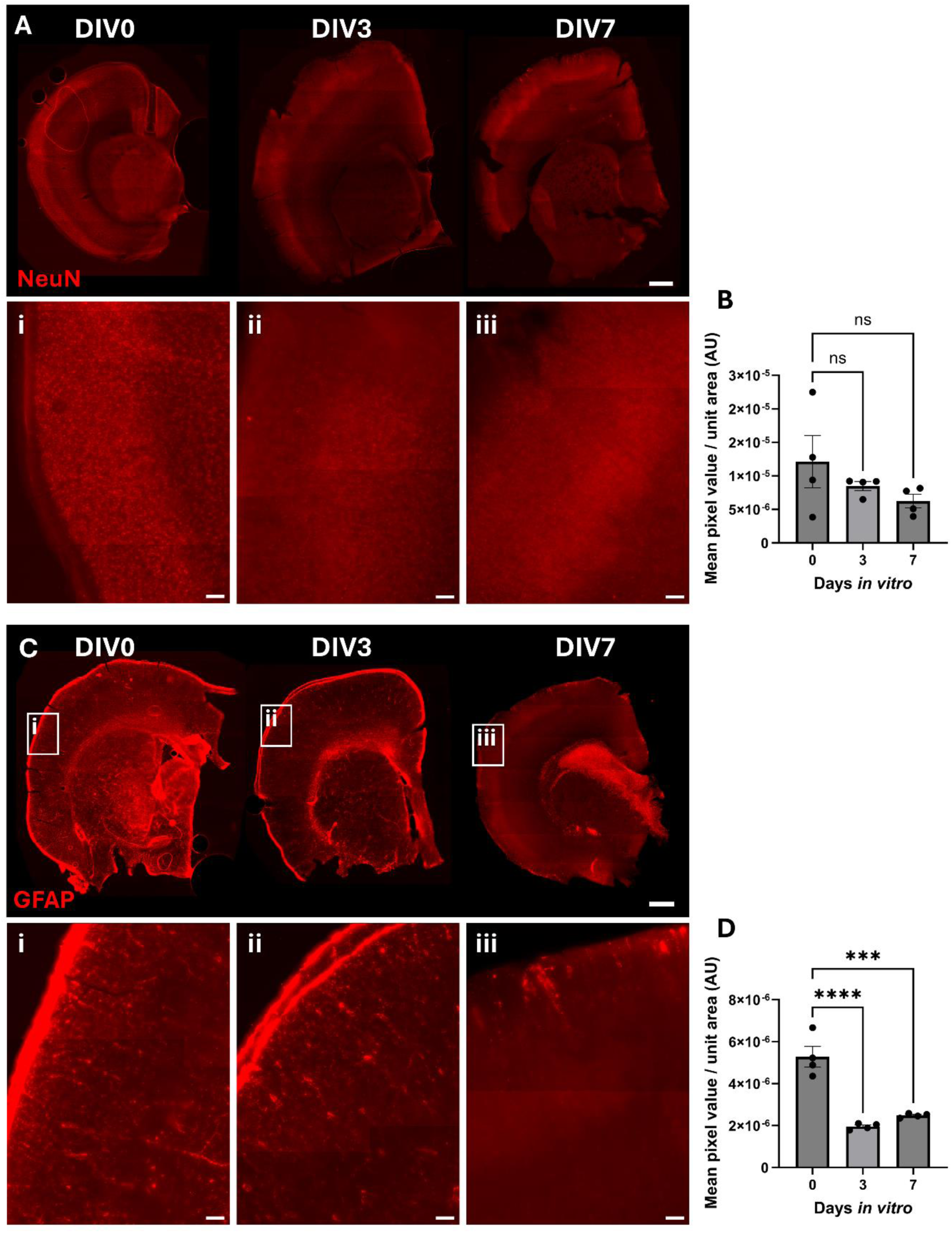
Immunofluorescent staining of neurons and astrocytes in OBSCs after 0, 3 and 7 days *in* vitro (DIV). Representative immunofluorescent staining of A) NeuN (neuronal cell marker) and C) GFAP (astrocyte marker) shown in red, with DAPI nuclear counterstain in blue (scale bars = 1,000 µm). Magnified regions of the cortex of section are shown in insets ‘i’, ‘ii’ and ‘iii’ for DIV0, 3 and 7 respectively (scale bars = 100 µm in insets), in addition to quantification of the mean pixel value per unit area for these regions in insets B) NeuN quantification in cortex, D) GFAP quantification in cortex. Each datapoint is the mean of 3 regions of interest per brain slice, ±SEM. All statistics were performed with one-way ANOVA with Dunnett’s multiple comparisons test (all to control). ns = not significant, ** *P* < 0.01, *** *P* < 0.001, **** *P* < 0.0001. *N* = 4 animals, *α* = 0.05.

Ionized calcium-binding adapter molecule 1 (IBA1) was used as a microglial cell marker. There was a reduction in IBA1-positive cells in the cortex in DIV3 and DIV7 compared to DIV0 (Figure 4A). Image analysis showed a significant reduction after 3 and 7 days of culture, from a mean IBA1 pixel intensity per unit area of 4.87×10^-6^ at DIV0 to 1.69×10^-6^ at DIV3 and 1.63×10^-6^ at DIV7 (Figure 4B, *P* = 0.0150 and 0.0134 respectively). However, similar to GFAP staining, there did not appear to be any significant changes in IBA1 staining in the subcortical white matter (Figure S4).

**Figure 4.**
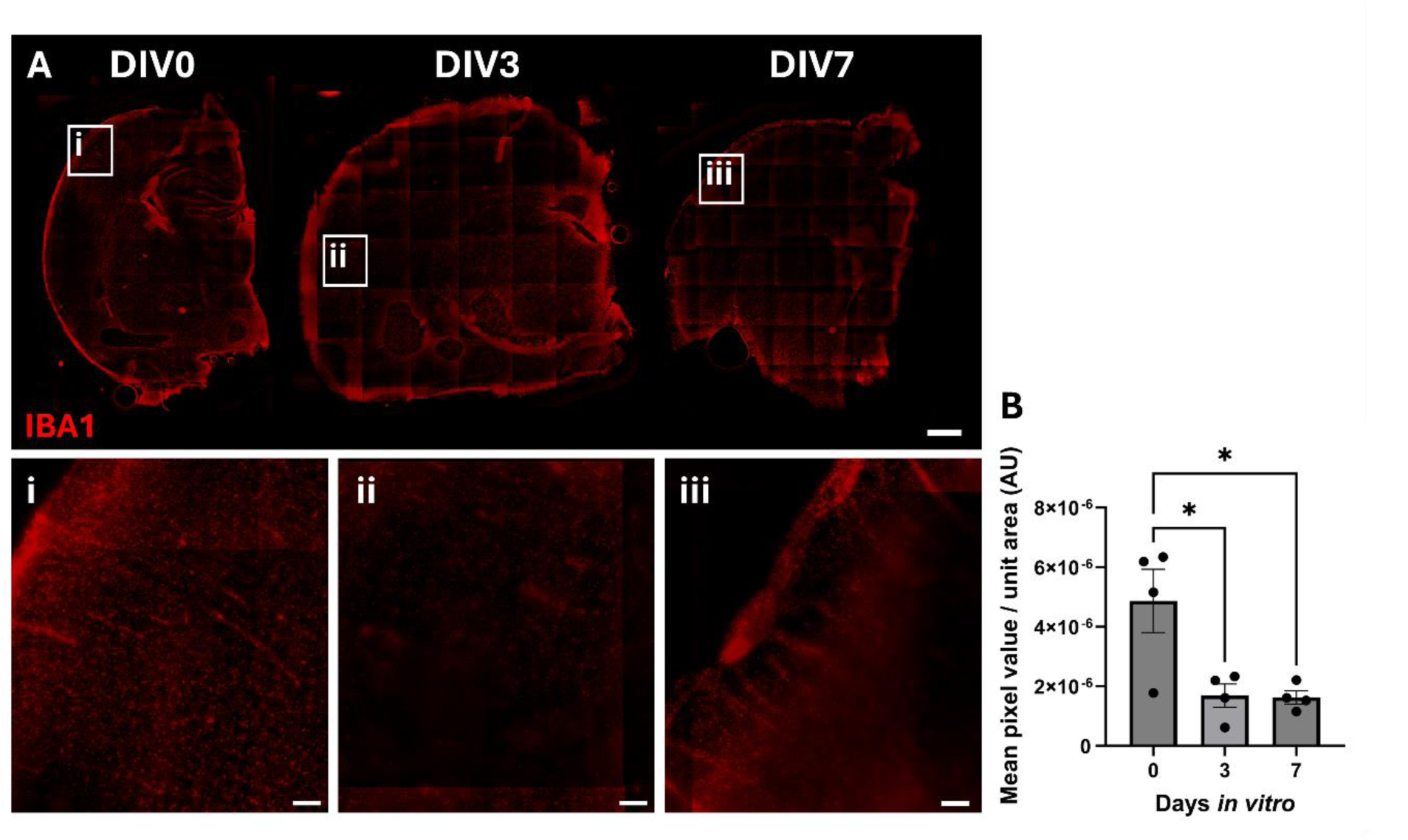
Immunofluorescent staining of microglia in OBSCs after 0, 3 and 7 days *in* vitro (DIV). Representative immunofluorescent staining of IBA1 shown in red (scale bars = 1000 µm). Magnified regions of the cortex of section are shown in insets ‘i’, ‘ii’ and ‘iii’ for DIV0, 3 and 7 respectively, in addition to quantification of the mean pixel value per unit area (scale bars = 100 µm in insets). Each datapoint is the mean of 3 regions of interest per brain slice, ±SEM. All statistics were performed with one-way ANOVA with Dunnett’s multiple comparisons test (all to control). ns = not significant, * *P* < 0.05. *N* = 4 animals, *α* = 0.05.

Alpha-smooth muscle actin (αSMA) labels vascular smooth muscle cells present within arteries, arterioles and pericytes. In contrast to the other cell markers, αSMA-positive staining (Figure 5A) increased along blood vessel structures in the cortex between DIV0 and DIV3 and DIV7. Image analysis revealed a significant increase in αSMA mean pixel value per unit area from 1.1×10^-6^ at DIV0 to 2.4×10^-6^ at DIV7 (*P* = 0.0061) (Figure 5B). Finally, rat endothelial cell antigen 1 (RECA-1) staining, which recognizes a cell surface antigen on endothelial cells, showed a reduction in staining of small vessels and capillaries at DIV3 and 7 (Figure 5C). Image analysis (Figure 5D) showed a marked and significant reduction in mean RECA-1 pixel value per unit area in the cortex, falling from 5.4×10^-6^ at DIV0 to 1.9×10^-6^ and 1.8×10^-6^ at DIV3 and DIV7 respectively (*P* = 0.0003 for both). A similar outcome was shown in the deeper tissues in proximity to the basal ganglia (Figure S4).

**Figure 5.**
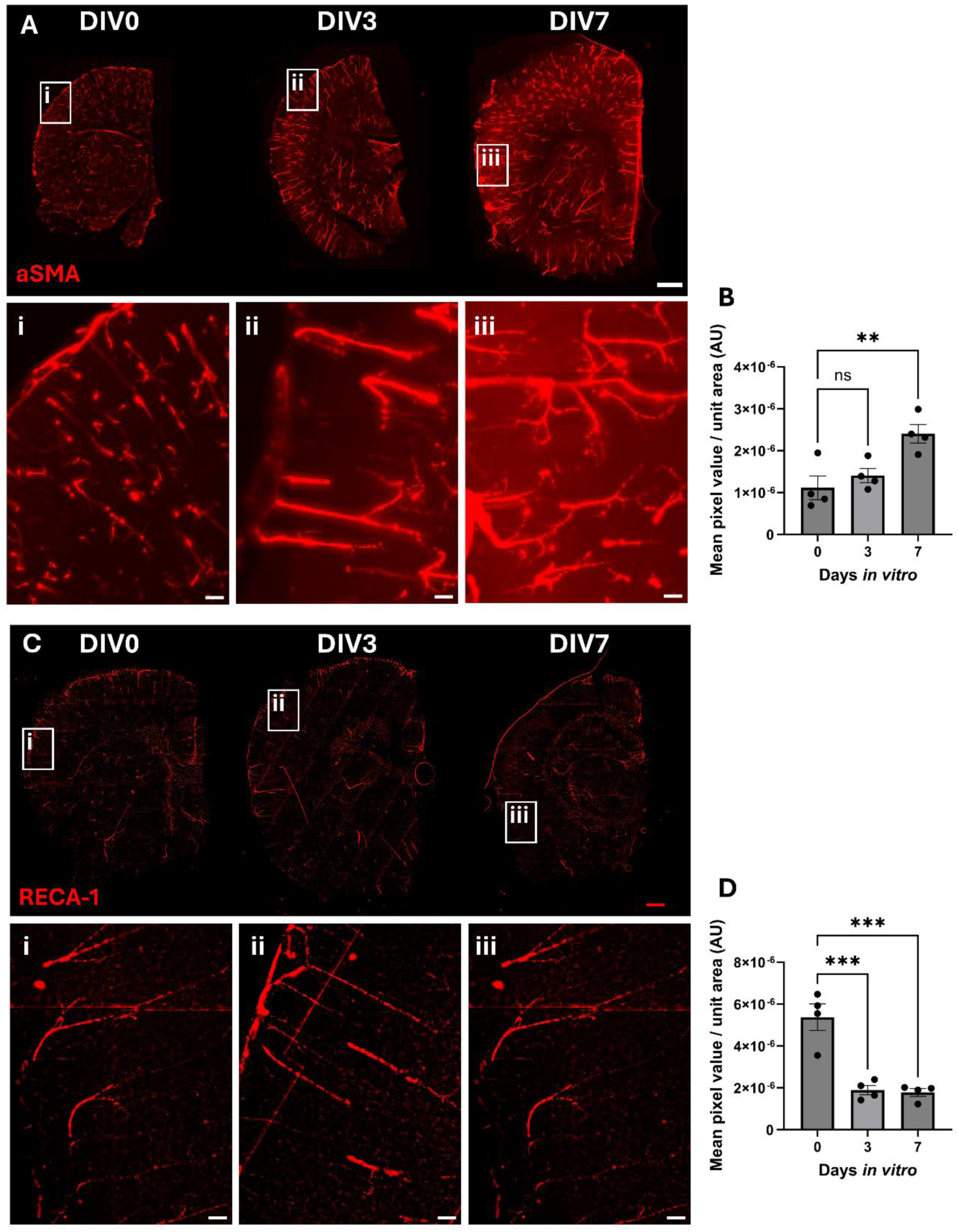
Immunofluorescent staining of smooth muscle cells, pericytes and endothelial cells in OBSCs after 0, 3 and 7 days *in* vitro (DIV). Representative immunofluorescent staining of A) αSMA (alpha smooth muscle actin) and C) rat endothelial cell antigen 1 (RECA-1) shown in red (scale bars = 1,000 µm). Magnified regions of the cortex of section are shown in insets ‘i’, ‘ii’ and ‘iii’ for DIV0, 3 and 7 respectively (scale bars = 100 µm in insets), in addition to quantification of the mean pixel value per unit area for these regions B) αSMA quantification, D) RECA1 quantification. Each datapoint is mean of 3 regions of interest per brain slice, ±SEM. All statistics were performed with one-way ANOVA with Dunnett’s multiple comparisons test (all to control). ns = not significant, ** *P* < 0.01, *** *P* < 0.001, **** *P* < 0.0001. *N* = 4 animals, *α* = 0.05.

### 4.3. Blood exposure induced increases in fibronectin and collagen deposits in OBSCs

OBSCs were exposed to lysed blood for 6, 24 or 48 hours to mimic haemorrhagic stroke, initiated at either DIV0 (acute) or DIV7 (7-day preculture). Fibronectin staining was shown to be minimal in acute sections (Figure 6A), with image analysis in the cortex and subcortical white matter (Figure 6B and Figure S8, respectively), and Western blotting of cortical protein isolates (Figure 6C and D) showing no significant increase in fibronectin staining or expression at any timepoint (*P* > 0.05 for all). However, control OBSCs cultured for 7 days showed visually greater fibronectin staining (Figure 6E) compared to acute sections (Figure 6A). Exposure of these pre-cultured OBSCs to blood resulted in a significant, near 10-fold increase in cortical vessels co-locating with fibronectin, from a mean of 2.97% in control OBSCs to a mean of 28.61% following blood exposure for 48 hours (*P* = 0.0123) (Figure 6F). This was further evidenced by a significant increase in fibronectin protein expression as assessed by Western blot, increasing 2.27-fold following blood exposure (*P* = 0.038) (Figure 6G and H). No significant change was noted in fibronectin staining in the subcortical white matter (Figure S8).

**Figure 6.**
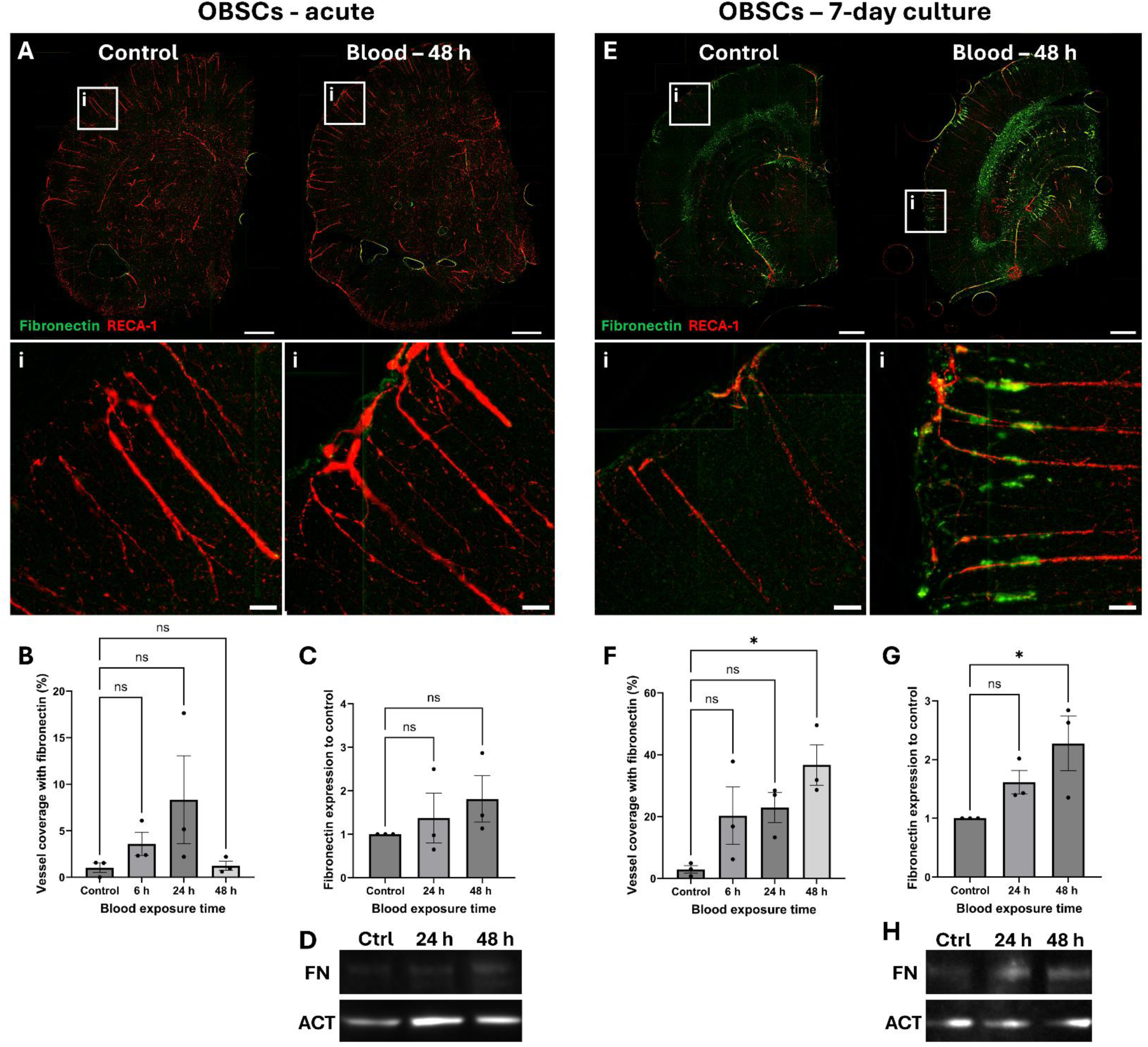
Localisation and quantification of fibronectin in blood-exposed OBSCs. OBSCs were generated and cultured either for 0 or 7 days, after which they were exposed either to PBS (control) or the same volume of lysed blood for 48 hours. OBSCs were then fixed and stained for RECA1 (red) and fibronectin (green). Representative images of whole hemispheres are shown (A – acute, E – 7 day pre-cultured, scale bars = 1,000 µm). Insets show magnified regions of the cortex (i) (scale bars = 100 µm). Quantification of the percentage coverage of fibronectin on RECA1-stained vessels was performed (B, F).

Fibronectin protein expression at 24 and 48 hours of blood exposure was also assessed by Western blot (D, H), followed by quantification of fibronectin fold-change (C, G). Each image quantification datapoint is the mean of three regions of interest from one brain slice per animal, ± SEM. All statistics were performed with one-way ANOVA with Dunnett’s multiple comparisons test (all to control). ns = not significant, *N* = 3 animals, *α* = 0.05.

A similar outcome was noted in the staining and expression of collagen IV, with acute OBSCs showing a significant increase in cortical vessel coverage from 1.55% in control samples to 16.15% after 24 hours of blood exposure (*P* = 0.0491, Figure 7B). No significant difference was seen in collagen IV staining in the white matter (*P* > 0.05 for all, Figure S8). In contrast to acute cultures, OBSCs treated with blood at DIV7 generated a marked increase in collagen IV signal, increasing from 7.92% cortical vessel coverage by collagen IV in the control to 32.84% after blood exposure (*P* = 0.0032, Figure 7D).

**Figure 7.**
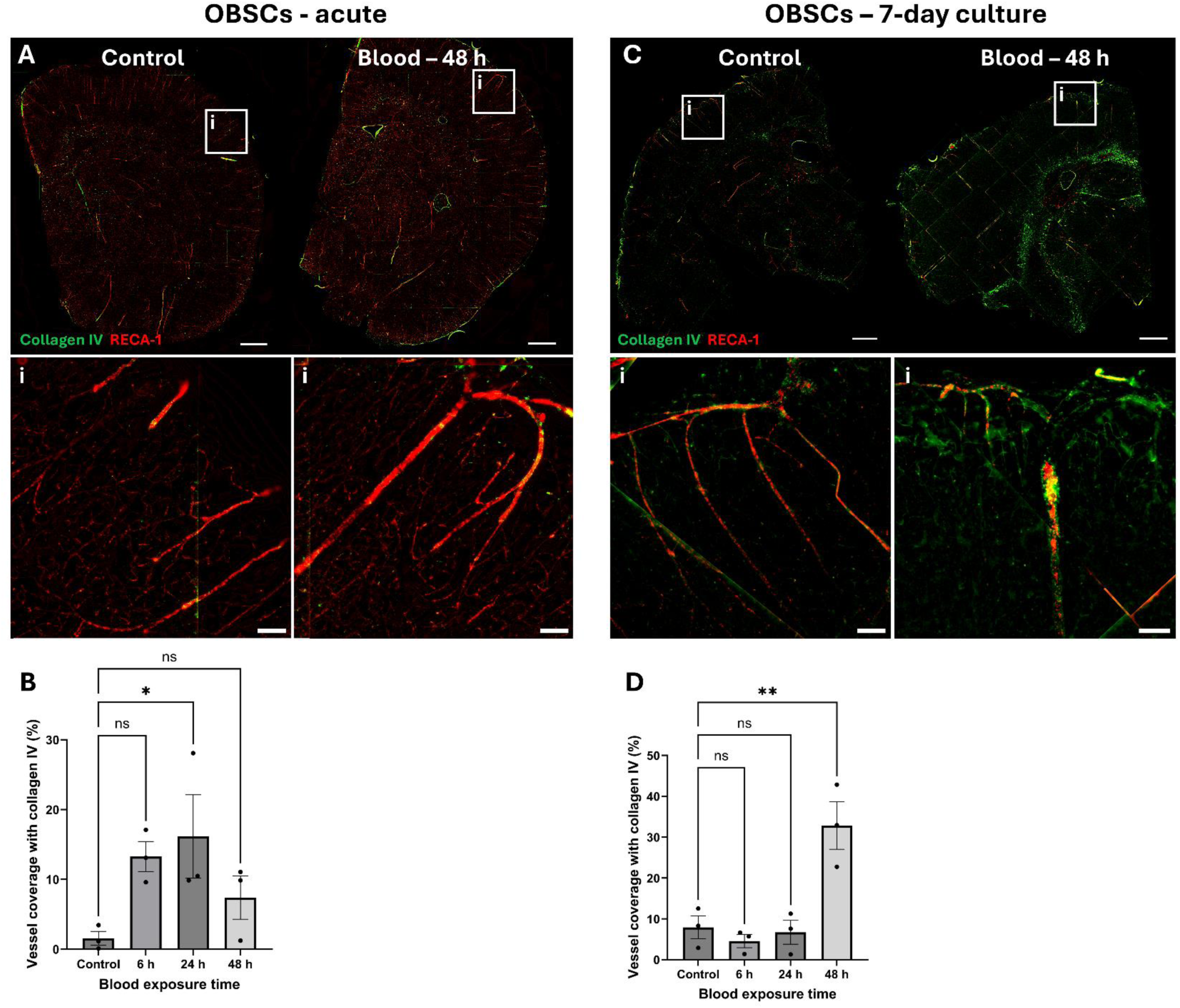
Localisation and quantification of collagen IV in blood-exposed OBSCs. OBSCs were generated and cultured either for 0 or 7 days, after which they were exposed either to PBS (control) or the same volume of lysed blood for 48 hours. OBSCs were then fixed and stained for RECA1 (red) and collagen IV (green). Representative images of whole hemispheres are shown (A – acute, C – 7 day pre-cultured, scale bars = 1,000 µm). Insets show magnified regions of the cortex (i) (scale bars = 100 µm). Quantification of the percentage coverage of collagen IV on RECA1-stained vessels was performed (B, D). Each image quantification datapoint is the mean of three regions of interest from one brain slice per animal, ± SEM. All statistics were performed with one-way ANOVA with Dunnett’s multiple comparisons test (all to control). ns = not significant, *N* = 3 animals, *α* = 0.05.

### 4.4. RNA-sequencing of blood-exposed OBSCs

Following sample quality control and principal component analysis (supplementary materials Figure S1 and S2), one blood-treated sample (B1) was excluded due to sequencing of an insufficient sample quality leading to poor mapping to the reference genome. All subsequent analysis was performed without this sample. Volcano plots were generated for all differentially expressed genes (DEGs) (*pAdj* < 0.05) with expression fold changes (FC) > 2, (Figure 8A) and greater than 1.5 (Figure 8B), with 95 and 604 DEGs over this FC cutoff, respectively. Heatmaps were also generated for each sample across the 25 most upregulated and 25 most downregulated DEGs (Figure 8C).

**Figure 8.**
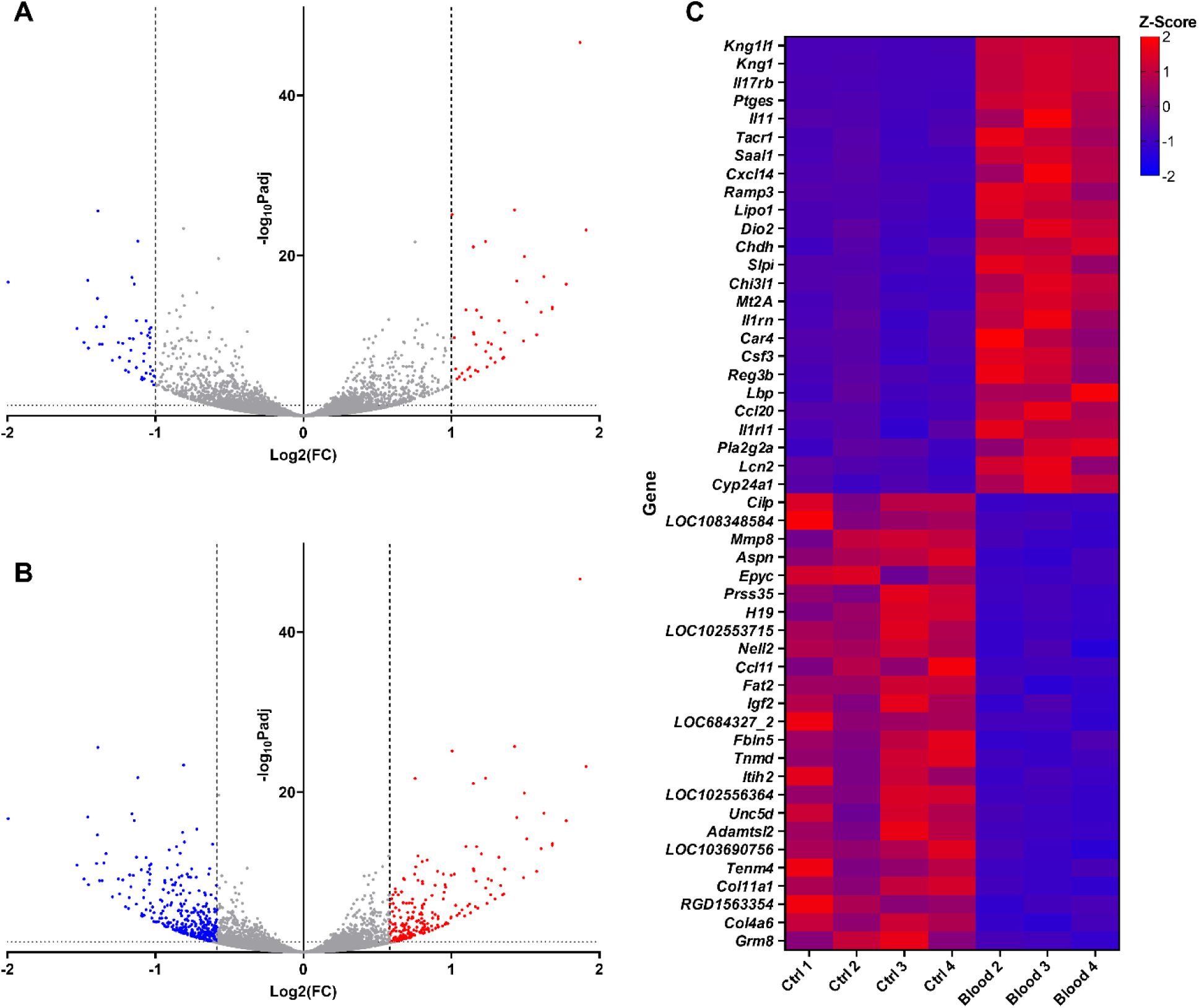
Volcano plots and heatmap of top 25 most differentially expressed genes (DEGs) in control and blood-treated OBSCs. Volcano plots set with a fold change threshold >2 (A) and >1.5 (B) (with fold change thresholds depicted by dotted vertical lines), showing upregulated DEGs (*pAdj* < 0.05) in red and downregulated DEGs in blue. Z-scores for top 25 most up- and down-regulated DEGs (ranked by log2 fold change) are visualised via heatmap (C), with upregulated genes shown in red and downregulated genes in blue.

RT-qPCR validation of the top 6 most upregulated and downregulated genes (Figure 9) showed a significant decrease in *Aspn*, *Epyc*, *H19*, *Mmp8* and *Prss35* transcript expression after blood exposure (*P* = 0.0107, 0.0159, 0.0027, 0.026 and 0.0297 respectively. There was no significant reduction in *Cilp* expression, likely due to high inter-sample variability (*P* = 0.2207). Inversely, a significant upregulation of *Il11*, *Ptges* and *Saal1* was shown (*P* = <0.0001, 0.0015 and 0.0006 respectively). Increases in *Kng1*, *Il17rb* and *Tacr* were not significant (*P* = 0.1307, 0.1109 and 0.0667). Overall, 8 of the 12 tested genes showed a significant change in expression (66.7%), with all showing a trend in the same direction as RNA-sequencing data following blood exposure.

**Figure 9.**
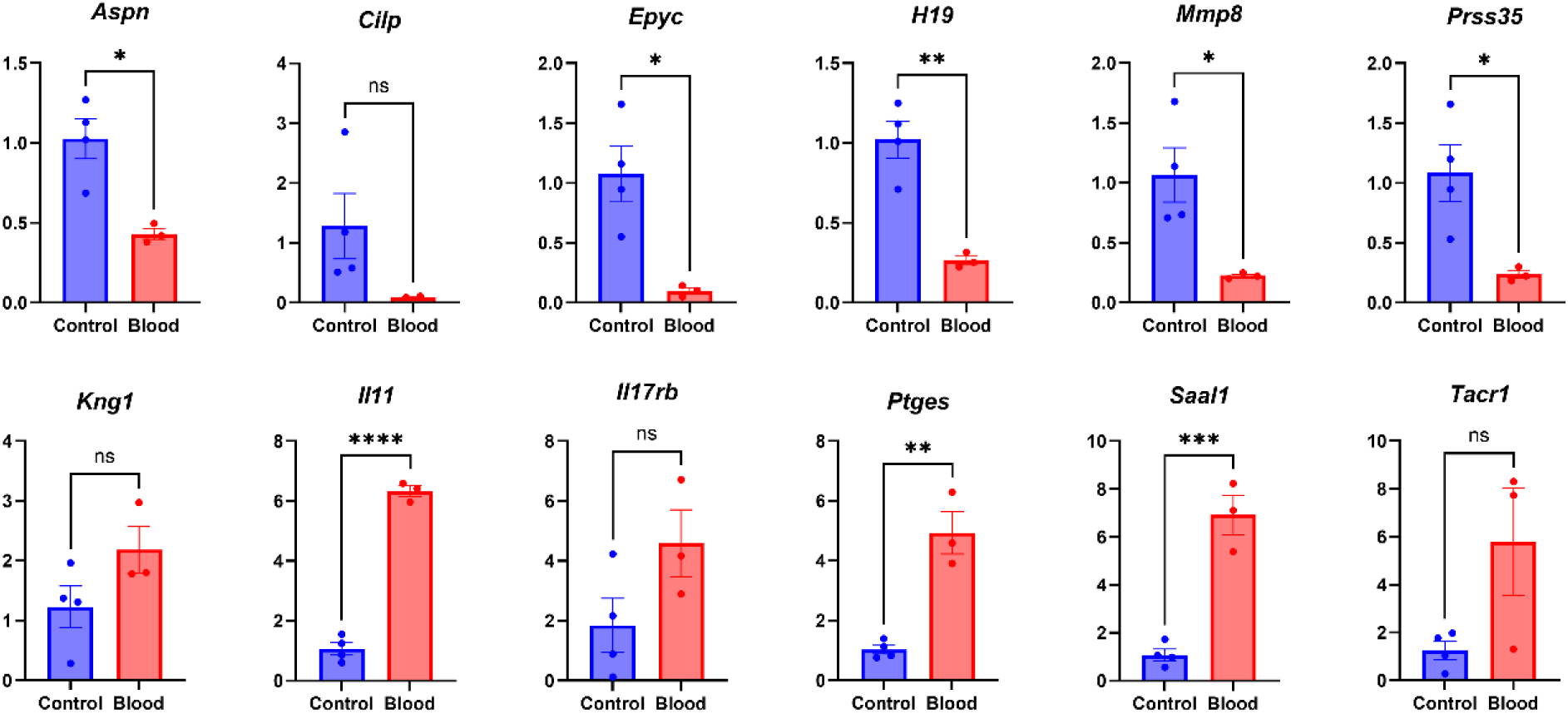
RT-qPCR validation of RNA-sequencing. Top 6 downregulated (lower expression in blood-exposed samples) and top 6 upregulated (higher expression in blood-exposed samples). All statistical analyses were performed with unpaired t-tests. ns = not significant, ns = not significant, * P < 0.05, ** P < 0.01, *** P < 0.001, **** P < 0.0001, *N* = 3, α = 0.05.

Gene ontology (GO) analysis was then performed across cellular component (CC), biological process (BP) and molecular function (MF) GO families. Within the CC family (Figure 10A), ‘extracellular region’ and ‘extracellular matrix’ were the top two most over-represented GO terms (gene ratios 0.137 and 0.285, *padj* 6.45x10^-29^ and 8.96x10^-27^ respectively). DEGs from extracellular region (Figure 10B) and extracellular matrix terms (Figure 10C) showed downregulation in blood-treated OBSCs including *Col11a1*, *Col4a6* and *Eln*, downregulation of *Mmp8* and upregulation of *Mmp9*, *Mmp3* and *Timp1*.

**Figure 10.**
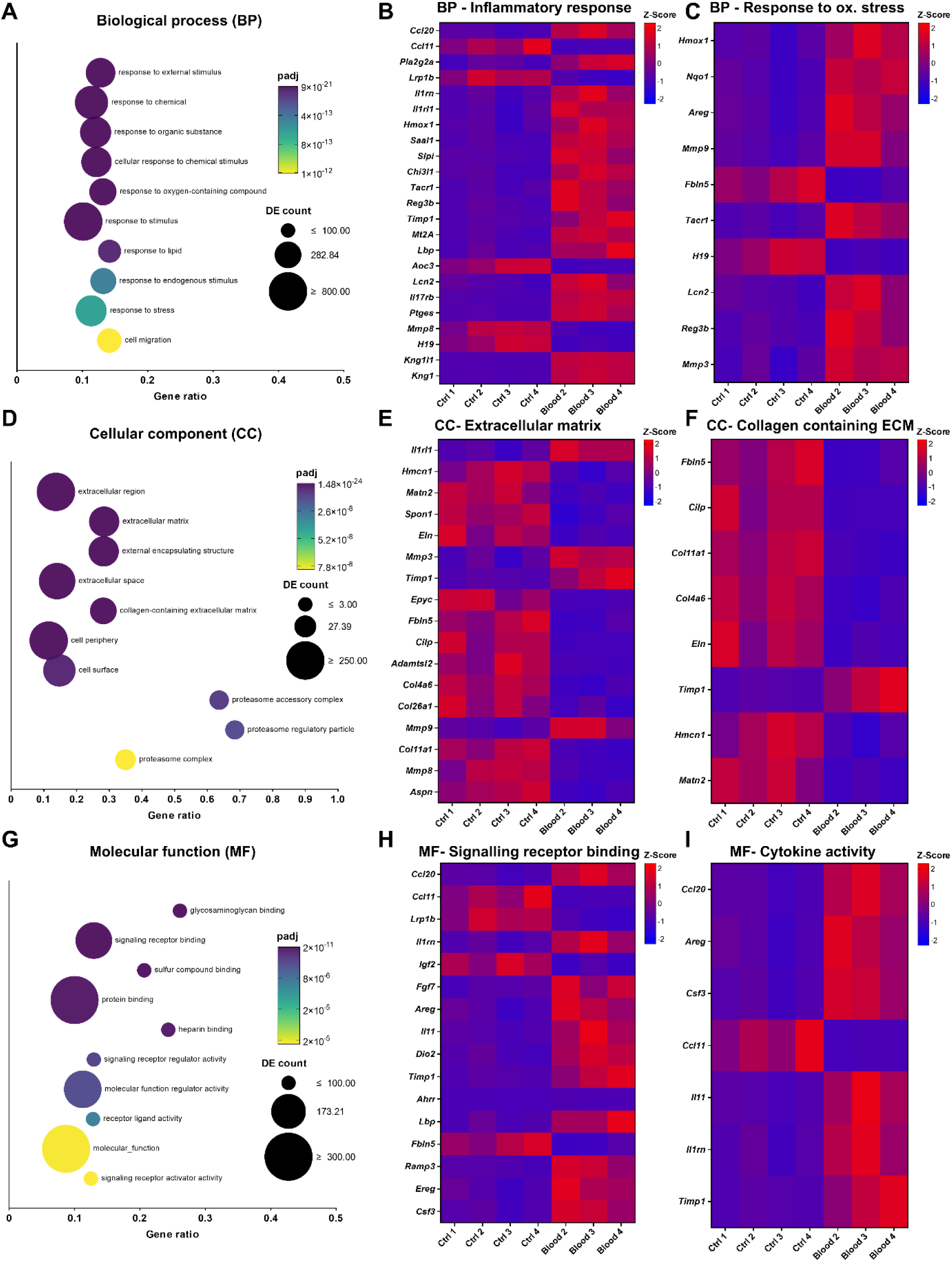
Gene ontology (GO) for control and blood-treated OBSC, with DEGs from selected GO terms within each family. A, D, G) Biological process, cellular component and molecular function GO families depicted with the top 10 most significant overrepresented CC GO terms shown as a bubble plot. The colour scale depicts adjusted p-value (*padj*) and bubble size represents the number of differentially expressed genes included in the respective category. Gene ratio of 1 = all genes in the GO category are DE, whilst 0 = none. B, C, E, F, H, I) Heatmaps of Z-score of DEGs with FC>2 or <0.5 within selected significantly over- represented GO terms ‘inflammatory response’, ‘response to oxidative stress’, ‘extracellular matrix’, ‘collagen containing extracellular matrix’, ‘signalling receptor binding’ and ‘cytokine activity’ respectively.

‘Response to external stimulus’ and ‘response to chemical’ were the top two most over- represented terms within the BP family (Figure 10D). ‘Inflammatory response’ and ‘response to oxidative stress’ were selected for display (gene ratios 0.141 and 0.149, *padj* 9.64x10^-11^ and 1.06x10^-7^ respectively), with DEGs from these terms shown in Figure 10E and F respectively. Blood exposure appeared to cause a marked upregulation of included cytokines and cytokine receptors, including *Ccl20*, *Il1rn*, *Il1r1* and *Il17rb*.

In the MF family, ‘glycosaminoglycan binding’ and ‘signalling receptor binding’ were the topmost over-represented terms (Figure 10G). ‘Signalling receptor binding’ and ‘cytokine activity’ were selected for display (gene ratios 0.129 and 0.168, *padj* 6.55x10^-10^ and 5.85x10^-5^), with DEGs from these terms shown in Figure 10H and Figure 10I, respectively. Blood exposure led to an increase in expression of *Il11*, *Csf3*, *Fgf7* and *Lbp* and a downregulation in *Igf2* and *Fbln5*, as examples.

## 5. Discussion

We have described an approach to the generation of adult rat OBSCs and their novel application to the development of an *ex vivo* haemorrhagic stroke model. We demonstrate that blood products have direct effects on ECM regulation in cortical blood vessels at both the transcript and protein level. In addition, we have characterised the distribution and number of several key brain cell types (neurons, endothelial cells, arterial smooth muscle/pericytes, astrocytes and microglia) after 0, 3 and 7 days in culture. All cell types were present across the measured time-course but showed differences in their response to culture conditions.

No significant change was noted in NeuN-positive cell staining, suggesting neurons were not substantially impacted by the length of time in culture. In contrast, there was a significant reduction in cortical GFAP-positive cells, particularly in the glia limitans, observed between DIV0 and DIV3/7. This could be due to either loss of astrocytes or reduction in GFAP expression within astrocytes. The latter seems plausible as the mechanical injury caused by vibratome sectioning has been shown to cause glial cell activation, a hallmark of which is GFAP upregulation ^23,24^. The reduction in GFAP levels observed in our study could therefore represent the return of GFAP to resting levels following an initial slicing-induced spike ^25^.

Quantification of astrocyte number is required to confirm this idea, but if correct, this interpretation would support the rationale for culturing brain slices for longer periods of time ^13^ before initiating studies in these models. Interestingly, GFAP levels were not reduced everywhere in the brain slice, and GFAP-positive staining was maintained over the time-course within the subcortical white matter. IBA1-positive microglia in the cortex also reduced from DIV0 to DIV3/7. Delbridge *et al*., reported a reduction in microglial proliferation and an increase in microglial inflammation in the days following slice preparation ^26^, likely leading to the observed fall in microglia numbers seen here.

We did observe a significant reduction in RECA-1 staining, particularly associated with capillaries, suggesting a loss of endothelial cells (ECs) from the OBSCs. We suggest that the lack of blood flow, and thus the lack of a pulsatile hemodynamic force and shear stress experienced by the ECs ^27^, contributes to this loss. Previous studies have shown that decreased blood flow directly affects EC migration and cytoskeleton reorganisation ^28^, and causes apoptosis and a reduction in EC number in previously dilated arteries ^29^. Alternatively, our data could be explained by a reduction in the abundance of the RECA-1 epitope (an unknown protein) within endothelial cells which has been suggested by Moser and colleagues^30^. At the same time, an increase in αSMA-positive cells was seen at DIV 7, suggesting a rise in the number of arterial smooth muscle cells (SMCs) or pericytes. Previous studies show a causative link between reduced blood flow and disordered vascular remodelling ^31^ and increased SMC proliferation (shown initially in hypertensive models) ^32^.

CCP3 staining increased between DIV0 and DIV3/7, indicating an increase in apoptosis-driven cell death, which is to be expected due to the acclimatisation of slices to culture following the injurious process of vibratome sectioning. This finding does suggest a possible upper limit to the culture period, though longer culture durations were not tested in the present work. Overall, these data suggest the successful maintenance of most tested cell types in OBSCs, with a possible degradation in blood vessel endothelium due to a lack of blood flow through the vasculature. For this reason, future works employing this model (for example, studies of the blood brain barrier) may choose to employ a shorter pre-culture period, with the caveat of increased astrogliosis and inflammation.

Following this, adult rat OBSCs were cultured for 0 and 7 days and then exposed to lysed rat blood for 6, 24 or 48 hours to mimic haemorrhagic stroke. Two key markers of fibrosis, fibronectin ^33,34^ and collagen IV ^35,36^,were both significantly elevated along cortical blood vessels (as measured by immunostaining) only in the 7-day cultures. 48 hours of blood exposure generated the largest response, both on immunofluorescent stains and Western blots. Our results show that these fibrotic deposits are primarily localised to penetrating cortical vessels, where αSMA is most prominent. Interestingly, these deposits were not observed in the 0-day cultures where there were lower levels of αSMA. This potentially indicating the involvement of SMCs or pericytes in modulating ECM associated with blood vessels. The location of ECM deposits suggest they could contribute to disordered glymphatic flow, reducing the clearance of metabolites (and blood products following haemorrhagic stroke) and exacerbating brain injury. Indeed, perivascular fibrosis has previously been reported in *in vivo* models of ICH ^37,38^ and SAH ^39^ and directly linked to reduced clearance of amyloid-β after ischemic stroke ^40^.

In addition to immunostaining and Western blotting, RNA sequencing was performed on RNA isolated from 7-day precultured OBSCs exposed to blood for 48 hours. Blood exposure led to significant over-expression of inflammatory response and cytokine signalling related GO terms, driven by a marked upregulation of inflammation-linked genes, including *Ccl20*, *Il11* and *Csf3*. *Ccl20* has previously been linked to neuroinflammation following traumatic brain injury ^41^ with antagonism shown to reduce this following SAH ^42^. *Il11* concentration in plasma post-ICH has been shown to positively correlate with increased mortality and the development of hydrocephalus ^43^. *Il11* has also been linked to inflammatory immune cell migration in multiple sclerosis ^44^ and protection against ischemia reperfusion injury in middle cerebral artery occlusion (MCAO) models of ischemic stroke ^45^. *Csf3* has been suggested to play a role in post-MCAO brain injury by increasing neutrophil toxicity ^46^. This large drive towards a pro-inflammatory transcriptome following blood exposure supports studies examining the pro-inflammatory nature of blood and blood products in the subarachnoid space and parenchyma following haemorrhagic stroke ^47,48^ and suggests that this crucial element is successfully recapitulated in our OBSC-based haemorrhagic stroke model.

Interestingly, the same RNA sequencing data showed a downregulation of ECM-related GO terms and genes (including collagens and elastin), with some upregulation of matrix metalloproteinases (MMPs) and tissue inhibitors of metalloproteinases (TIMPs).

Immunofluorescent staining and Western blotting of blood-exposed OBSCs suggested a marked increase in ECM deposition along cortical vessels, as previously discussed, suggesting a differential response at the gene and protein level. We propose this increase in protein deposition may not be due to changes in *de novo* protein synthesis, but a manipulation of ECM turnover triggered by blood-induced changes in the expression of MMPs and TIMPs. Indeed, modified *Mmp9*/*Timp1* expression (as shown in our RNA sequencing data) has been linked to disordered collagen deposition in the skin ^49^ and lung ^50^. In the brain, a reduction in collagen IV RNA and upregulation of *Mmp9* has previously been reported 48 hours after SAH ^51^, a finding reflected in the present work. These findings suggest that our model also successfully mimics the ECM dysfunction seen in haemorrhagic stroke, with a likely breakdown of the blood brain barrier and disordered, fibrotic ECM deposition along cortical penetrating vessels.

A limitation of our model is that it simulates global blood exposure instead of localised blood exposure to the cortical tissues or ventricles (as may be seen after SAH or IVH). Further work may be undertaken to identify if this model can recapitulate the differences seen between types of haemorrhagic stroke, i.e SAH and ICH ^8^. Single cell RNA sequencing may also determine which cells drive fibrosis following blood exposure, as the mechanism underlying this has not been explored in the present work.

Our findings suggest that our OBSC model can a) successfully maintain adult rat brain slices for at least 7 days and b) employ lysed blood to recapitulate the disordered ECM deposition (fibrosis) and neuroinflammation which follows haemorrhagic stroke. Future studies may employ this model to further understand the molecular drivers of post-stroke inflammation and fibrosis, or to trial anti-fibrotic agents to reduce ECM dysfunction. Our model offers a platform to advance the translation of pharmacological treatments aimed at reducing post-stroke disability and improving outcomes for stroke survivors, whilst providing a higher throughput and physiologically realistic alternative to *in vivo* animal studies.

## 6. Declarations

### 6.3. Data availability

RNA sequencing data have been uploaded to the NCBI sequence read archive (SRA) and are freely available for use (BioProject ID PRJNA1262501). Uncropped, unedited Western blots are included in the supplemental materials (Figures S3 and S4). Image quantification macros are available (supplemental materials, ‘ImageJ macro language scripts used for image quantification’). All other data will be made available upon request to the corresponding authors.

### 6.4. Authors’ contributions (CRediT)

BJH: methodology, software, formal analysis, resources, investigation, data curation, writing – original draft, visualisation. LR: investigation, data curation, writing – review and editing. JAR: formal analysis, writing – review and editing. DF: resources, writing – review and editing. PK: supervision, writing – review and editing. RMB: supervision, writing – review and editing. LJH: supervision, writing – review and editing. HB: resources, writing – review and editing, supervision, project administration, funding acquisition.

## 7. Acknowledgements

The authors thank the members of the Neuro-Ocular Inflammation and Matrix Research Group (University of Birmingham) for their support and University of Birmingham Biomedical Services Unit staff for their expert assistance in caring for the experimental animals. We acknowledge the support of the Microscopy Technology Hub Facilities at the College of Medicine and Health, University of Birmingham, for providing access to equipment and technical expertise and the Birmingham Environment for Academic Research (BEAR) advanced research computing team for use of high-performance computing services and storage.

The Galaxy server used for some calculations is partly funded by the German Federal Ministry of Education and Research BMBF grant 031 A538A de.NBI-RBC and the Ministry of Science, Research and the Arts Baden-Württemberg (MWK) within the framework of LIBIS/de.NBI Freiburg. The Aston Institute for Membrane Excellence (AIME) is funded by UKRI’s Research England as part of their Expanding Excellence in England (E3) fund.

