## Supplemental materials file 1 for "Adult organotypic brain slice cultures recapitulate extracellular matrix remodelling in haemorrhagic stroke"

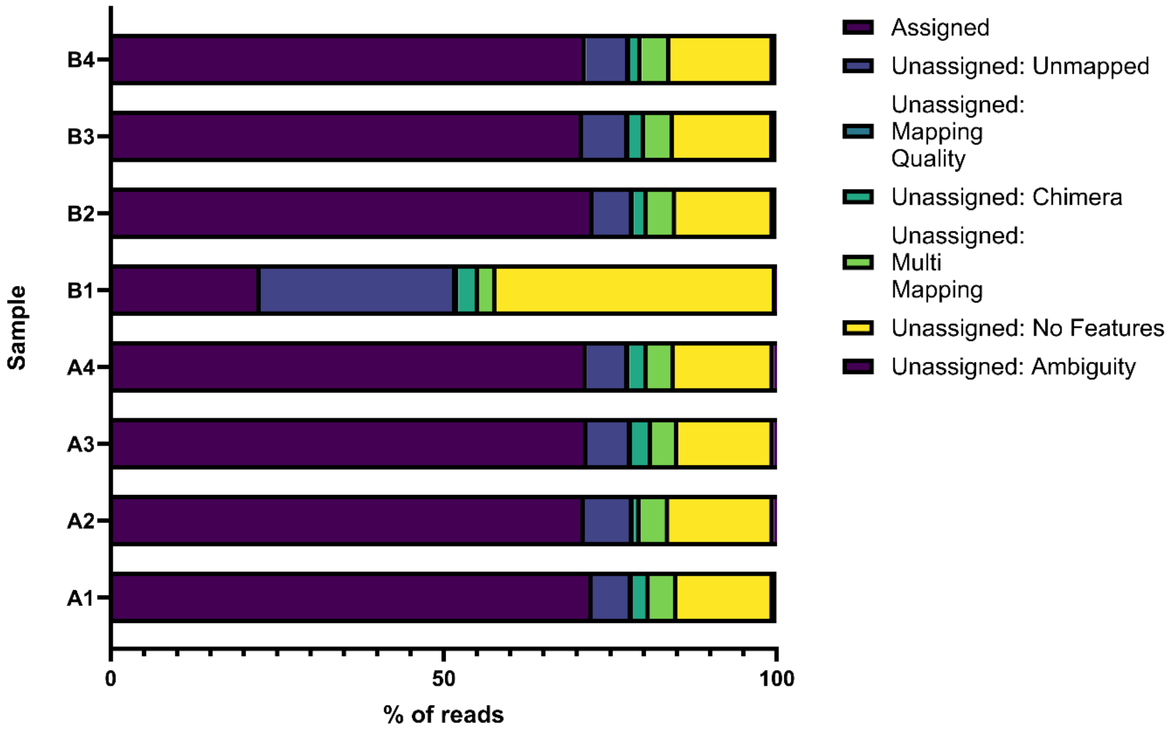


**Figure S1 - Mapping of RNA-seq reads to reference genome**

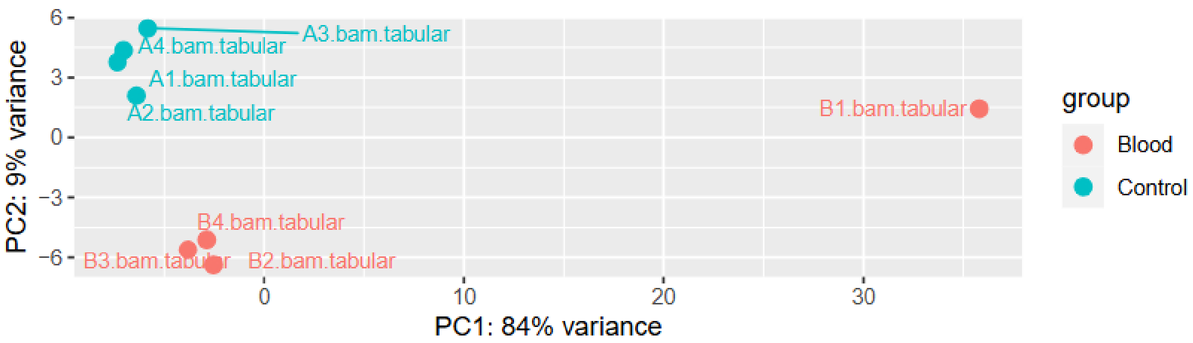


**Figure S2 - Principal component analysis (PCA) of control (blue) and blood-exposed (red) OBSC RNA**

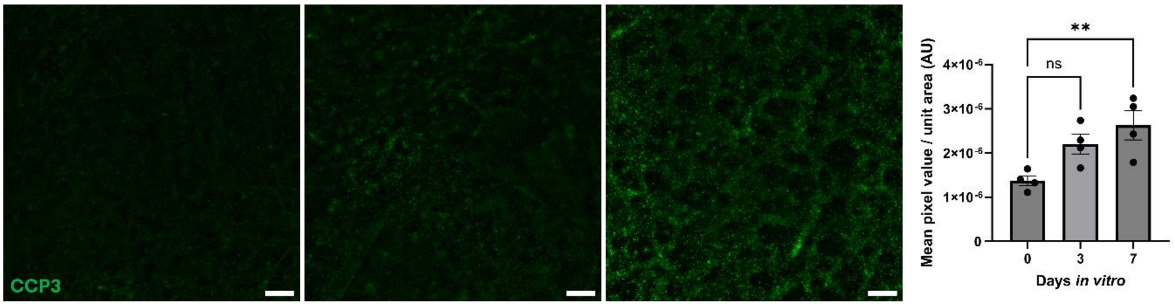


**Figure S3 - Distribution and intensity of CCP3 throughout basal ganglia of control and cultured OBSCs.** A) Representative immunofluorescent staining of OBSCs (CCP3, green and DAPI, blue) at 0, 3 and 7 days *in vitro* (scale bar = 1000 µm). Magnified regions of the cortex are shown in insert i (scale bar = 100 µm for all). Images processed by background subtraction (50 px radius) and enhancement of window and level for all images equally to improve visibility. Quantification of mean CCP3-stained pixel intensity per unit area in the basal ganglia was also performed (B). Each image quantification datapoint is mean of 6 regions of interest from 2 brain slices per animal (3 regions per slice), ± SEM. One-way ANOVA with Dunnett's multiple comparisons test (all to all). ns = not significant, ** $P < 0.01$, *** $P < 0.001$. $N = 4$ animals, $\alpha = 0.05$.

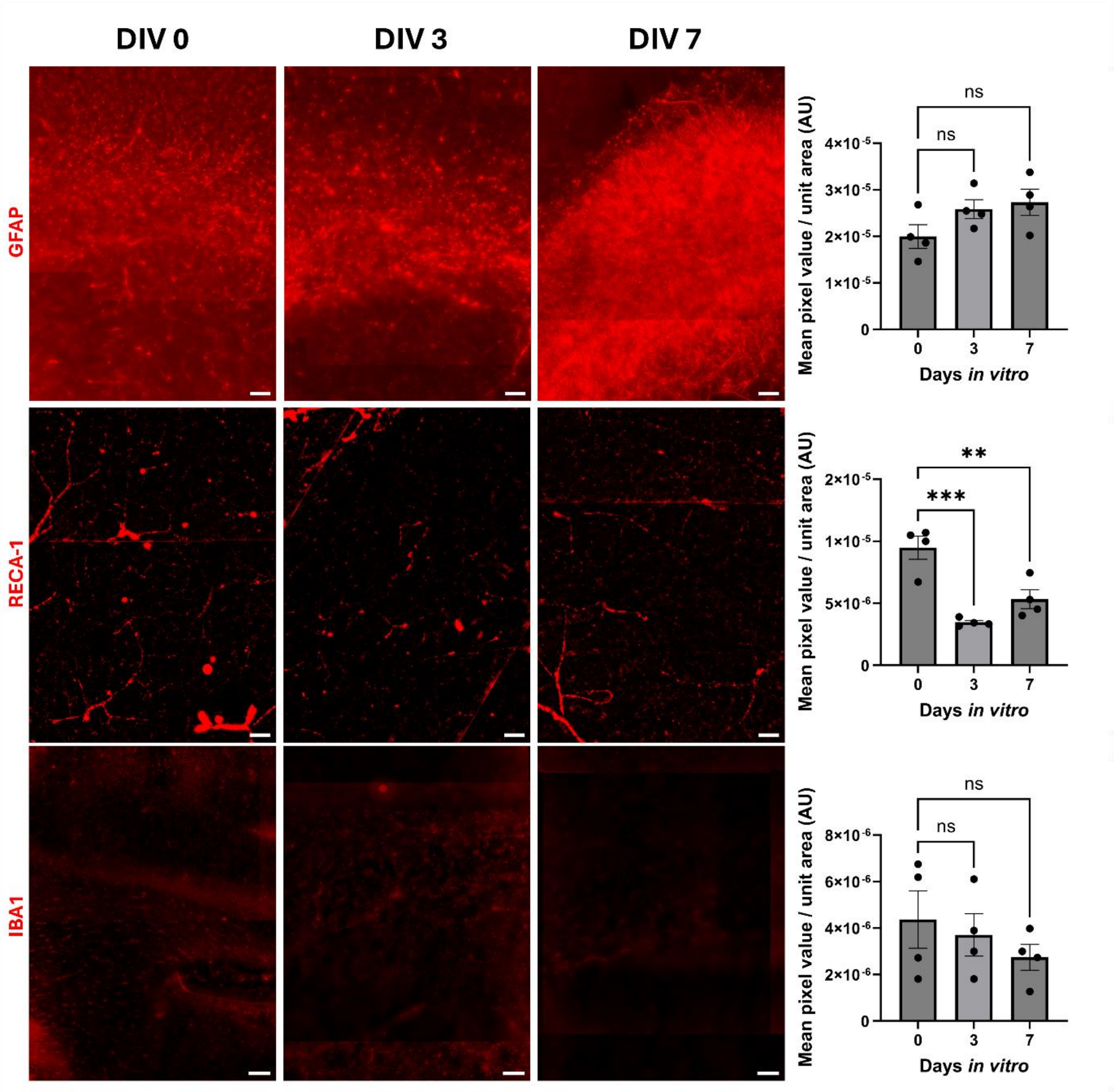


**Figure S4 - Immunofluorescent staining of astrocytes, blood vessels and microglia in deeper tissues of OBSCs after 0, 3 and 7 days in vitro (DIV).** Regions of interest are sub-cortical white matter for GFAP and IBA1, basal ganglia for RECA-1. Scale bars 100 µm. Quantification of mean stained pixel intensity per unit area was also performed. Each image quantification datapoint is mean of three regions of interest from one brain slice per animal ± SEM. One-way ANOVA with Dunnett's multiple comparisons test (all to control). ns = not significant, ** $P < 0.01$, *** $P < 0.001$. $N = 4$ animals, $\alpha = 0.05$.

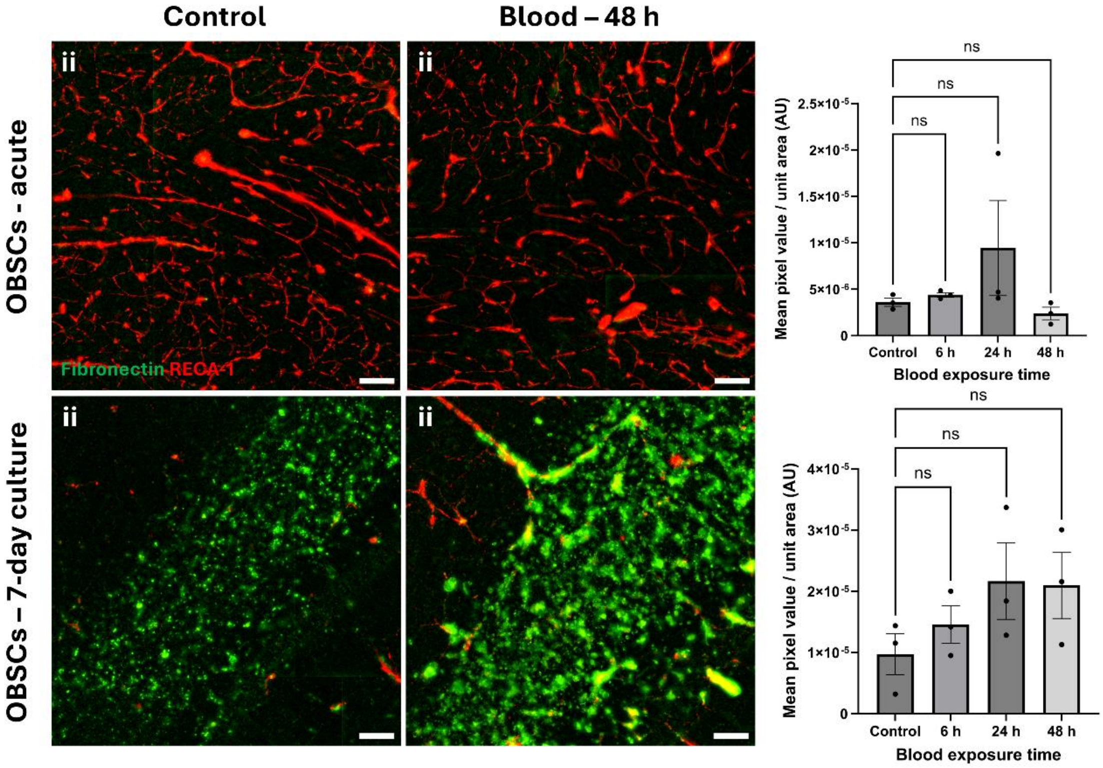


**Figure S5 – Localisation and quantification of fibronectin in sub-cortical white matter of blood-exposed OBSCs.** OBSCs were generated and cultured either for 0 or 7 days, after which they were exposed either to PBS (control) or the same volume of lysed blood for 48 hours. OBSCs were then fixed and stained for RECA1 (red) and fibronectin (green). Scale bars = 100 µm. Quantification of the percentage coverage of fibronectin on RECA1-stained vessels was performed (B, D). Each image quantification datapoint is mean of three regions of interest from one brain slice per animal, ± SEM. All statistics performed with one-way ANOVA with Dunnett's multiple comparisons test (all to control). ns = not significant, N = 3 animals, $\alpha = 0.05$.

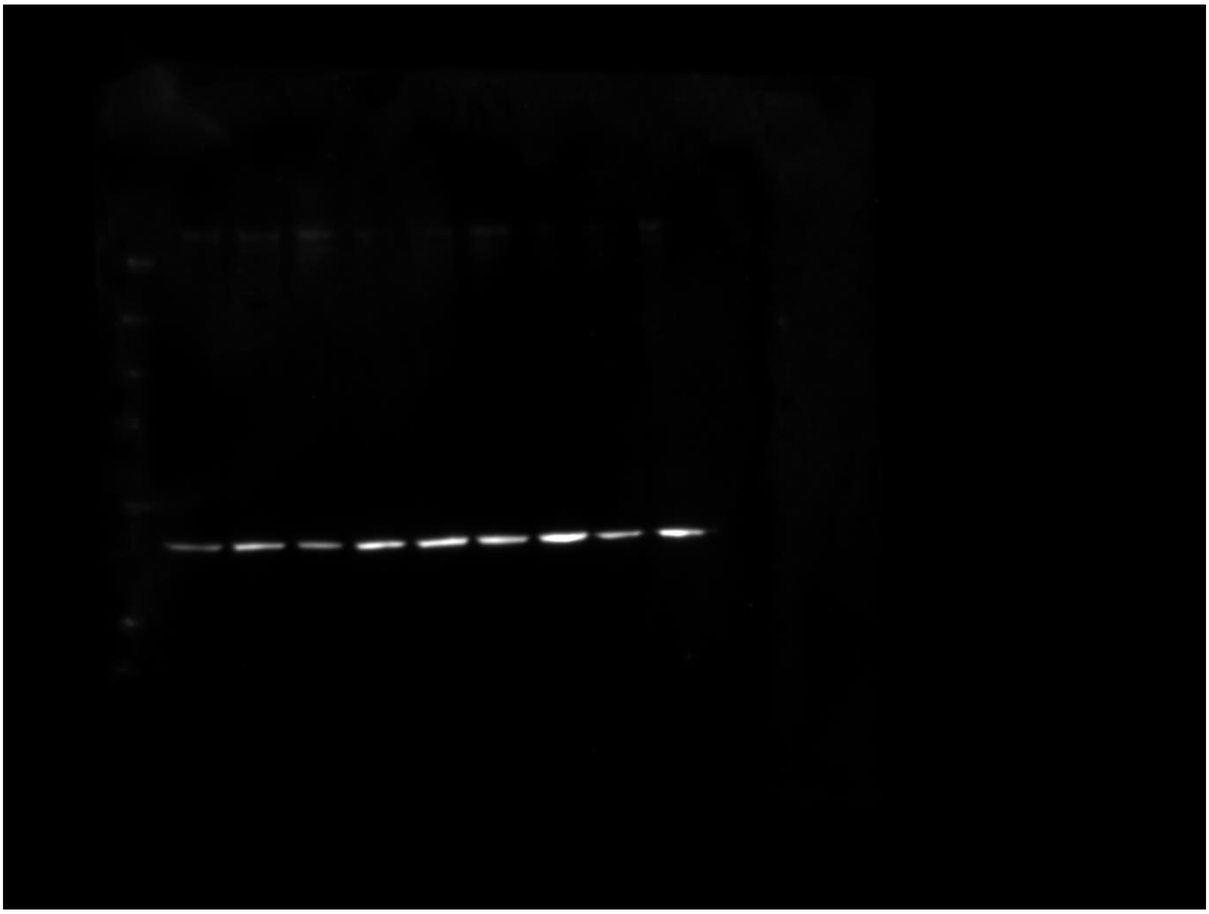

**Figure S6 - Unedited, uncropped western blot for acute blood exposed OBSCs.** Lanes (L-R)– Ladder, N1 Ctrl, N1 24hrs blood, N1 48hrs blood, N2 ctrl, N2 24 hrs blood, N2 48hrs blood, N3 ctrl, N3 24 hrs blood, N3 48 hrs blood.

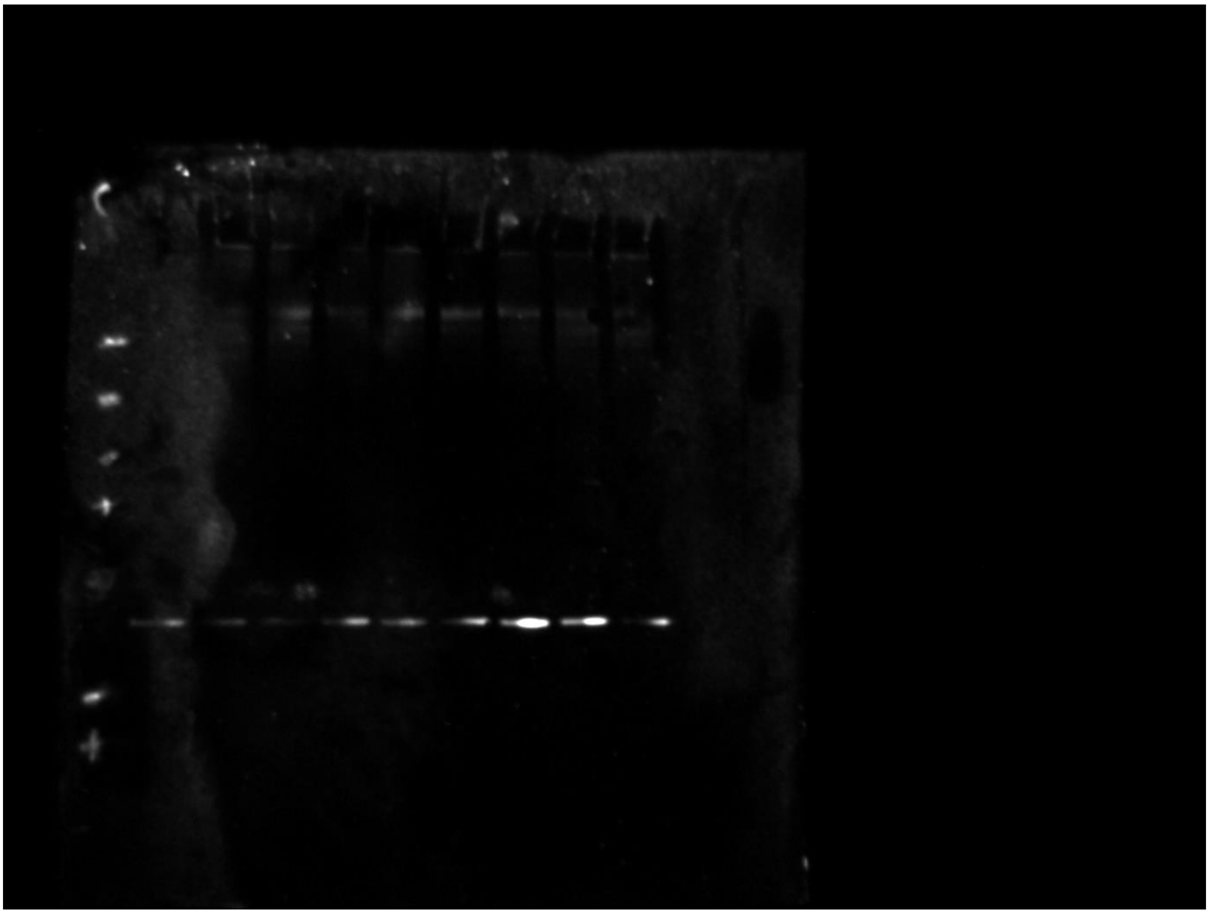

**Figure S7 - Unedited, uncropped western blot for 7-day precultured blood exposed OBSCs.** Lanes – Ladder, N1 Ctrl, N1 24hrs blood, N1 48hrs blood, N2 ctrl, N2 24 hrs blood, N2 48hrs blood, N3 ctrl, N3 24 hrs blood, N3 48 hrs blood.

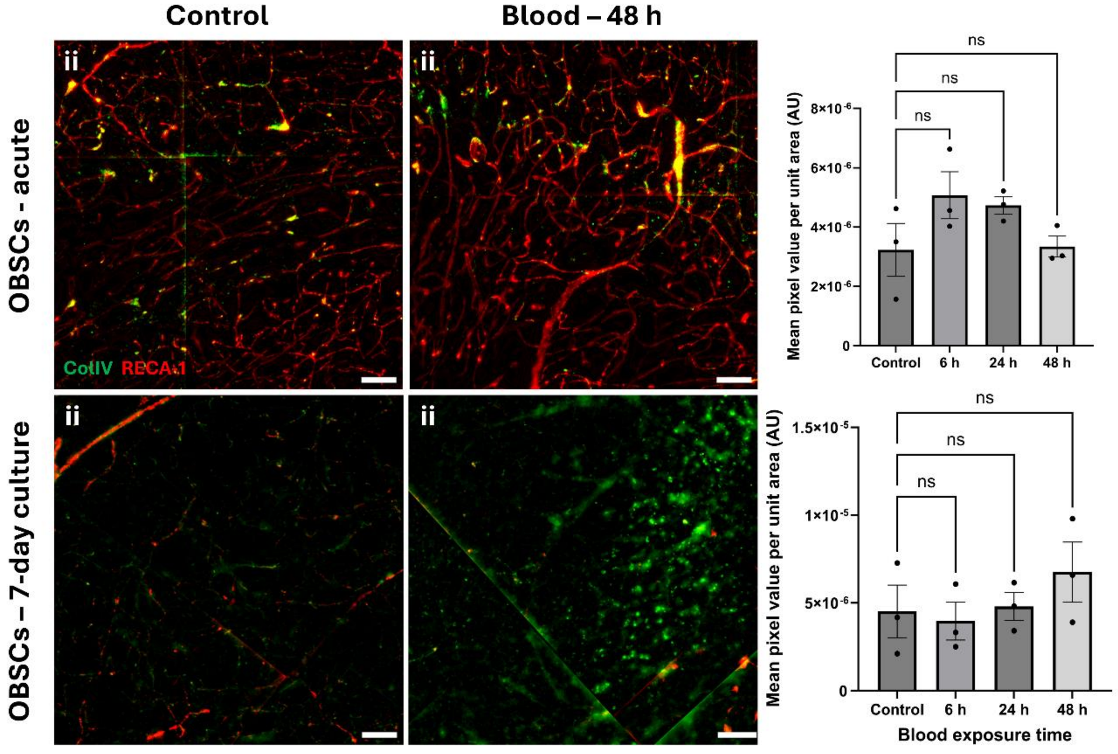


**Figure S8 - Localisation and quantification of collagen IV in sub-cortical white matter of blood-exposed OBSCs.** OBSCs were generated and cultured either for 0 or 7 days, after which they were exposed either to PBS (control) or the same volume of lysed blood for 48 hours. OBSCs were then fixed and stained for RECA1 (red) and collagen IV (green). Scale bars = 100 µm. Quantification of the percentage coverage of collagen IV on RECA1-stained vessels was performed (B, D). Each image quantification datapoint is mean of three regions of interest from one brain slice per animal, ± SEM. All statistics performed with one-way ANOVA with Dunnett's multiple comparisons test (all to control). ns = not significant, N = 3 animals, $\alpha = 0.05$.

### RT-qPCR validation of RNA-sequencing

**Table S1 - RT-qPCR reaction conditions for RNAseq validation.**

| Reaction step | Temperature (°C) | Ramp rate (°C/s) | Hold time (mm:ss) | Cycles |
|---|---|---|---|---|
| 1 – Preincubation | 95 | 4.4 | 05:00 | 1 |
| 2 - Amplification | - | - | - | 45 |
| 2a | 90 | 4.4 | 00:10 | - |
| 2b | 60 | 2.2 | 00:10 | - |
| 2c | 72 | 4.4 | 00:10 | - |
| 3 – Melting Curve | - | - | - | 1 |
| 3a | 95 | 4.4 | 00:05 | - |
| 3b | 65 | 2.2 | 01:00 | - |
| 3c – Acquisition (5 acquisitions / °C) | 65-97 | 0.11 | - | Continuous |
| 4 - Cooling | 4 | 2.2 | 30:00 | 1 |

**Table *S*2 - Oligonucleotide primers used in RT-qPCR validation of RNAseq.**

| Gene symbol | Transcript | Forward (5'-3') | Reverse (5'-3') |
|---|---|---|---|
| *KNG1* | ENSRNOT00000078131.2 | CCAGGGTGCAAGAAGAGAGG | CCCATGCTTATGACCTTGGTG |
| *IL17RB* | ENSRNOT00000020823.7 | GCATTCATACAGCTGCGTGA | GGGAAGCCGACATAGGAGAA |
| *PTGES* | ENSRNOT00055011957.1 | GAGGTCTCCAGTACTGCAGG | GATCGTCTCCATGTCGTTGC |
| *IL11* | ENSRNOT00000023489.6 | CTTCAGACCCTCGTGCAGAT | GCCAAGGTAGGTAGGGAGTC |
| *TACR1* | ENSRNOT00000007984.6 | GCTGGCCATGAGTTCTACCA | TCCAGCCCCTCATAATCACC |
| *SAAL1* | ENSRNOT00000016254.6 | CAAATACTTCCATGCTCGGGG | CAGCTCTTGAGTCCTCTGCT |
| *CILP* | ENSRNOT00000044887.5 | AAGGTCACCCAACTCACTGT | GGTGGCATTCTGGAAGCAAT |
| *MMP8* | ENSRNOT00000013936.4 | CATAGCAAGCGTGTTCCCAG | AGTGACTCTGCGACTGACAA |
| *ASPN* | ENSRNOT00000074041.3 | ACGTGTGAGAGAGATCCACT | ATCTTTGGCACTGTTGGACA |
| *EPYC* | ENSRNOT00000006450.8 | GCAACAACAGACTCGGAAGG | TGGTCCAAGCTGTTATCCGT |
| *PRSS35* | ENSRNOT00000034683.5 | AGGACACAGCAAGCTTCTCA | TGGTTCCGTTTCAGATCCCA |
| *H19* | ENSRNOT00000102917.1 | GCTGCTCTCTGGATCCTCTT | GAAGTCCCCGGATTCAAAGG |
| *ACTB* | ENSRNOT00000080216.2 | TCAGGTCATCACTATCGGCA | AGGTCTTTACGGATGTCAACG |
| *RPL27* | ENSRNOT00000028060.7 | ACTACAACCACCTCATGCCC | TCCCTGTCTTGTATCGCTCC |
| *RPL13A* | ENSRNOT00000093559.2 | TGCAAAAGCTTCTGGAGGATG | AGGTGGGAGATGTTGGTCTG |
| *YWHAZ* | ENSRNOT00000080676.2 | ATGATGGGAGGCAGTGAGTC | TAATTCTCCTCAGCCTGGGA |

### ImageJ macro language scripts used for image quantification

Also available at - https://github.com/Dr-ben-hewitt/OBSC-quantification-macros

**Percentage coverage of blood vessels:**

```
// Define the input and output directories
inputDir = getDirectory("Choose the input directory containing CZI files");
interDir = getDirectory("Choose the intermediate file directory (just needs an empty folder)");
ROIoutputDir = getDirectory("Choose a folder for the ROIs");
outputDir = getDirectory("Choose the output directory for TIFF files");



fileList = getFileList(inputDir);
firstImage = true;
windowMin = 0;
windowMax = 255;
// Loop through each file in the input directory
for (i = 0; i < fileList.length; i++) {
  // Check if the file has a .czi extension
  if (endsWith(fileList[i], ".czi")) {
    // Open the CZI file
    open(inputDir + fileList[i]);
```

```
    // Get the title of the current image (used to name the output file)
    title = getTitle();
    origTitle = getTitle();
    noExt = File.nameWithoutExtension;
            if (firstImage) {
    // Wait for the user to adjust the window and level manually
            waitForUser("Open the W/L window, adjust, and click apply. Then click OK.");

    // Get the current window and level values
            getMinAndMax(windowMin, windowMax);

    // Set the firstImage flag to false
             firstImage = false;
             }
            else {
    // Apply the saved window and level values to subsequent images
              setMinAndMax(windowMin, windowMax);
            }
        run("Z Project...", "projection=[Max Intensity]");
            run("8-bit");
            Stack.setChannel(1);
    run("Green");
    Stack.setChannel(2);
    run("Red");
            saveAs("OME-TIFF", interDir + "MIP_" + noExt);
            close(); //added
  }
}
run("Close All");
interFileList = getFileList(interDir);
for (x = 0; x < interFileList.length; x++) {
  // Open the current image
```

```
    open(interDir + interFileList[x]);
    // Wait for the user to select ROIs
    Stack.setChannel(2);
    waitForUser("Please select ROIs from image as required, pressing 't' to add them to the
manager");
    // Get the number of ROIs
    n = roiManager("count");
    roinum = 1;
    // Loop through all ROIs in the ROI Manager
    for (i = 0; i < n; i++) {
        // Get the current image title and remove extension
        title = getTitle();
        noExt = File.nameWithoutExtension;
        roiManager("Select", i);
        run("Duplicate...", "duplicate");
        saveName = noExt + "_" + "ROI-" + roinum;
        saveAs("OME-TIFF", ROIoutputDir + saveName);
        close();
        roinum++;
    }
    roiManager("Deselect");
    roiManager("Delete");
}
run("Close All");
////////Split, background subtract, merge ////////////////////////////////////////////
outputFileList = getFileList(ROIoutputDir);
textDir = getDirectory("Choose the histogram output directory");
textFile = textDir + "output.csv";
File.append("Image Title, Histogram Values\n", textFile);
for (x = 0; x < outputFileList.length; x++) {
                open(ROIoutputDir + outputFileList[x]);
        title = getTitle();
        origtitle = getTitle();
```

```
    noExt = File.nameWithoutExtension;
            imageSeq = 0;
run("Split Channels");
// PROCESS CH1///////////
selectWindow("C" + 1 + "-" + title);
run("Subtract Background...", "rolling=1000");
            run("Auto Threshold", "method=RenyiEntropy white");
//PROCESS CH2 ////////////////
selectWindow("C" + 2 + "-" + title);
run("Subtract Background...", "rolling=100");
run("Auto Threshold", "method=Triangle white");
ch1 = "C1-" + title;
ch2 = "C2-" + title;
imageCalculator("Average create", ch1, ch2);
//run("Merge Channels...", "c1=" + ch1 + " c2=" + ch2 + " create");
merge_title = "Processed_" + noExt;
rename(merge_title);
print("Finished " + merge_title);
selectWindow(merge_title);
            //// Histogram section
            getHistogram(values, counts, 256);
            line = noExt;
            for (j = 0; j < 256; j++) {
    line = line + counts[j];
    if (j < 255) {
    line = line + ",";
            }
    }
    line = line + "\n";
    File.append(line, textFile);
selectWindow(merge_title);
            saveAs("OME-TIFF", outputDir + merge_title);
```

```
    // Close all images
    run("Close All");
 }

print("Processing complete!");
```

**Intensity per unit area:**

```
// Define the input and output directories
inputDir = getDirectory("Choose the directory containing the images to be analysed (TIFF format)");
outputDir = getDirectory("Choose a folder for the output");
//
inputFileList = getFileList(inputDir);
textDir = getDirectory("Select the histogram output directory");
textFile = textDir + "GreenQuant.csv";
File.append("Image Title, mean pixel value, area \n", textFile);
for (x = 0; x < inputFileList.length; x++) {
   // Open the current image
   open(inputDir + inputFileList[x]);
   title = getTitle();
   noExt= File.nameWithoutExtension;
   print(x);
   run("Split Channels");
   selectWindow("C1-" + title);
   run("Subtract Background...", "rolling=50 stack");
   run("Measure");
   meanvalue = getResult("Mean");
   areavalue = getResult("Area");
   saveName = "GREEN-BGSub-50__" + title;
         line = saveName + ","+ meanvalue + "," + areavalue + "\n";
   File.append(line, textFile);
   saveAs("OME-TIFF", outputDir + saveName);
```

```
  //

          run("Close All");

}

print ("Quantification done");
```

### Staining and clearing of OBSCs

1) Heparin solution – 10 mg Heparin (10KU) to 1ml PBS, aliquot into 125 µl, store at -20°C
2) PBS+tritonX100 (2% v/v) – 49 ml PBS + 1 ml Triton X
3) 50% methanol in PBS – 25 ml methanol to 25 ml PBS
4) 80% methanol in dH2O – 40 ml methanol to 10 ml dH2O
5) 100% methanol
6) 20% DMSO in methanol – 40 ml methanol to 10 ml DMSO
7) Hydrogen peroxide (5% v/v) in methanol-DMSO (80-20% v/v) – Base of 8 ml DMSO to 32 ml methanol, add 8 ml 30% H2O2
8) Antibody penetration buffer (APB) – PBS with 0.2% triton-X and 20% DMSO + 0.3M glycine (MW = 75.07 g/mol). *1.6 ml per sample.*
    a. 39.9 ml PBS
    b. 10 ml DMSO
    c. 0.1 ml Triton-X100
    d. 1.126 g glycine
9) Blocking buffer (BB) – PBS with 0.2% triton, 6% goat serum and 10% DMSO. *0.8 ml per sample.*
    a. 41.9 ml PBS
    b. 5 ml DMSO
    c. 0.1 ml TritonX
    d. Store until needed, and then add 0.6 ml goat serum per 9.4 ml BB needed
    e. Store at 4°C with serum added, maximum of 3 months
10) Antibody dilution buffer (ADB) – PBS with 0.2% tween20, 10 µg/ml heparin, 3% serum, 5% DMSO. *0.8 ml per sample.*
    a. 45.85 ml PBS
    b. 2.5 ml DMSO
    c. 0.1 ml Tween20
    d. Store until needed, and then add 0.3 ml serum and 10 µl of 10 mg/ml heparin to 9.69 ml
    e. Store at 4°C with heparin added, maximum of 3 months
11) Wash buffer – PBS with 0.2% Tween20 and 10 µg/ml Heparin. *1.6 ml per sample.*
    a. 499 ml PBS
    b. 1 ml Tween20
    c. Store until needed, then add 100 µl of 10 mg/ml heparin per 99.9 ml
    d. Store at 4°C with heparin added, maximum of 3 months

1) Except where otherwise stated, perform all steps in the procedure at room temperature with gentle agitation.

1) Fix brain slices in 4% PFA for 60 mins at room temperature.

2) Wash tissues in PBS for 60 mins x 2
3) Permeabilize tissues by washing them in increasing concentrations of methanol at 4°C with gentle agitation.
    a) PBS for 8 mins x 2
    b) 50% methanol in PBS for 8 mins
    c) 80% methanol in deionized water for 8 mins
    d) 100% methanol for 8 mins
4) Bleach tissues by submerging in ice-cold 5% $H_2O_2$ in 20% DMSO/methanol, then incubating at 4°C overnight.
5) Wash samples in 1.6mls of:
    a) 20% DMSO/methanol for 8 mins
    b) 80% methanol in deionized water for 8 mins
    c) 50% methanol in PBS for 8 mins
    d) 100% PBS for 8 mins
    e) PBS with 2% Triton™ X-100 for 8 mins
6) Incubate the samples in 1.6ml Antibody Penetration Buffer for 30 mins with gentle shaking.
7) Block the samples in 0.8ml Blocking Buffer for 60 mins with gentle shaking at 37°C.
8) Transfer the samples to 0.2ml primary antibody prepared in Antibody Dilution Buffer then incubate at 37°C with gentle shaking for 90 mins.
9) Wash the samples in 1.6ml Wash Buffer for 20 mins with gentle shaking x 5
10) Incubate the samples in 0.2ml secondary antibody prepared in Antibody Dilution Buffer at 37°C with gentle shaking for 90 mins.
11) Wash the samples in 1.6ml Wash Buffer at 4°C overnight
12) Wash samples in 1.6 ml Wash Buffer at 37°C with gentle shaking for 20 mins x 9
13) Dehydrate the tissues with increasing concentrations of ethanol (used due to incompatibility of CellBrite with methanol) at 4°C with gentle shaking. Wash tissues in 2mls of:
    a) 50% methanol in PBS for 8 mins
    b) 80% methanol in deionized water for 8 mins
    c) 100% methanol for 8 mins
14) Remove the tissues from methanol, scooping onto labelled slide and ensure that all excess methanol is absorbed
15) Add 1ml CytoVista™ Tissue Clearing Reagent, then incubate at 4°C with gentle shaking for 20 mins
16) Mount on slides in CytoVista™ Tissue Clearing Reagent.
17) Coverslip and seal with nail polish
